# Quantifying seed rain patterns in a remnant and a chronosequence of restored tallgrass prairies in north central Missouri

**DOI:** 10.1101/2024.07.10.602969

**Authors:** K. Carter Wynne, Maya J. Parker-Smith, Erica M. Murdock, Lauren L. Sullivan

## Abstract

1. Seed rain is an influential process related to plant community diversity, composition, and regeneration. However, knowledge of seed rain patterns is limited to those observed in forests and late-assembling grasslands, which might not reflect early-assembling communities such as newly restored grasslands. Resolving this gap in our understanding provides further insight into the role of seed dispersal.
2. Here, we measured seed rain in a remnant tallgrass prairie, which was the site of the foundational grassland seed rain study in 1978, and a nearby chronosequence of tallgrass prairie restorations. We sought to determine how the quantity, seed mass traits, timing, diversity, and composition of seed rain changed (1) long-term and (2) during community assembly. To do so, we deployed artificial turf grass seed traps into 2-year-old, 5-6-year-old, and 15-year-old restored prairies and the remnant prairie, replacing traps every two weeks from May to December 2019.
3. We captured over twice the density and richness of seed rain in the remnant prairie in 2019 compared to 1978. We also found that seed rain patterns changed as prairies aged, with each prairie possessing a distinct community of dispersing species. Significantly more seeds, seed biomass, and species were captured in the youngest restored prairie. However, seed mass traits were similar in all prairies. Except for composition, all other seed rain metrics in the oldest restoration were eventually comparable to the remnant prairie.
4. *Synthesis and Applications*: Our results revealed that grasslands, notably young prairies, produce larger quantities of seed rain than previously known (124,806 seeds m^−2^ year ^−1^, 97.24 g m^−2^ year ^−1^), and seed input in all sampled prairies far exceeded restoration broadcast seeding densities. We further found that decreases in seed rain quantity across the chronosequence did not correspond with increases in seed mass, suggesting a lack of tradeoffs between these metrics. Furthermore, tallgrass prairie restorations have not replicated the composition of seed rain seen in remnant systems. Increasing restoration seeding rates of desirable species may be needed to meet composition goals since current rates may not compete with the propagule pressure of undesirable species found in newly restored prairies.

## Introduction

Seed production and dispersal are critical processes of plant community dynamics (Beckman & Sullivan, 2023; Levine & Murrell, 2003). The dispersing pool of seeds, known as seed rain, represents an essential transition between plant generations produced via sexual reproduction (Arruda et al., 2018; Rabinowitz & Rapp, 1980). Seed rain plays a significant role in the regenerative ability (e.g., Huanca Nuñez et al., 2021; Schott & Hamburg, 1997), successional dynamics (e.g., Sullivan et al., 2018; Tilman, 1994), introductions of novel species (e.g., D’Antonio et al., 2001; DiVittorio et al., 2007), and maintenance of genetic and species diversity in plant communities (Beckman & Sullivan, 2023; Clark et al., 2007; Myers & Harms, 2009; Turnbull et al., 2000). Inadequate densities of dispersing propagules in the seed rain can limit plant population size and constrain local species diversity, a widespread phenomenon known as seed limitation (Clark et al., 2007; Foster, 2001; Grman et al., 2015; Martin & Wilsey, 2006; Myers & Harms, 2009; Turnbull et al., 2000). Even though seed dispersal is central to many processes related to plant community dynamics, studies on community-level seed rain remain rare (Beckman & Sullivan, 2023; Levine & Murrell, 2003).

Dispersal is key to overcoming seed limitation across ecosystems, particularly in disturbed, newly assembling communities (Clark et al., 2007; Myers & Harms, 2009; Turnbull et al., 2000). In many ecosystems, increasing the supply of dispersing propagules is often enough to overcome the negative effects of seed limitation and increase local species richness (Foster, 2001; Myers & Harms, 2009). For example, ecosystem restoration efforts use this principle to increase species diversity by seeding larger quantities of desirable species into restored communities (Rowe, 2010). However, pinpointing the role of seed input for population and community dynamics can be difficult when essential characteristics of ambient seed rain (e.g., quantity, diversity, and composition) are unknown (Clark et al., 2007; Myers & Harms, 2009). Despite its importance to all plant communities across community assembly, patterns of seed rain have primarily been studied in forests and mature grasslands (Arruda et al. 2018). As a result, experiments and restorations are often informed using seed rain estimates from late-assembling, mature systems, even though restoration efforts occur in early-assembling systems such as newly restored grasslands (e.g., Martin & Wilsey, 2006; Rowe, 2010). A lack of robust seed rain estimates for these systems makes it difficult to determine whether seeding densities are saturating enough to overcome seed limitation (Clark et al., 2007; Myers & Harms, 2009). Overall, resolving the existing mismatch in our understanding of seed rain at early-stages vs. late-stages of community assembly will allow for a better evaluation of the effects of seed dispersal.

Shifts in species traits relating to seed production, dispersal, and competitive ability during community assembly likely influence vital aspects of seed rain as communities age. Across ecosystems, life history and seed size are correlated with seed rain density, where fast-reproducing, short-lived, small-seeded species produce and disperse larger quantities of seeds than long-lived, large-seeded species that tend to delay reproduction (Moles et al., 2004; Moles & Westoby, 2006; Sullivan et al., 2018). The numerical advantage afforded by these smaller-seeded species is thought to be offset by large-seeded species having increased juvenile survival and competitive ability (Moles et al., 2004; Tilman, 1994; Turnbull et al., 1999). However, tradeoffs in seed production, dispersal, and competitive ability may differ in strength between ecosystems. For example, seed rain studies using a chronosequence approach in forest ecosystems observe seed rain dispersing into mature forests is more species-rich, contains more large-seeded species, and is compositionally different compared to early-assembling forest seed rain; however, seed rain quantity may increase (Piotto et al., 2019), decrease (Young et al., 1987), or not change (Huanca Nuñez et al., 2021) with age. Limited studies in grassland ecosystems suggest declines in seed rain input but increases in large-seeded, perennial species and diversity over time, supporting tradeoffs in seed rain density and seed mass (Kettenring & Galatowitsch, 2011; West & Durham, 1991). Seed rain in grasslands is also far denser than in forests, and abiotic rather than biotic vectors play more prominent roles in grassland seed dispersal (Kettenring & Galatowitsch, 2011; Rabinowitz & Rapp, 1980; Schott & Hamburg, 1997; West & Durham, 1991; Willand et al., 2015). Key aspects of seed rain may respond differently during community assembly in grasslands compared to forests. Therefore, measuring a suite of metrics related to seed rain is needed to critically assess how seed rain patterns change during grassland community assembly and whether these shifts are associated with changes in species traits.

To deepen our understanding of the role of dispersal during community assembly, we measured seed rain patterns in a remnant tallgrass prairie and a nearby chronosequence of restored prairies, ranging from 2 to 15 years old in Missouri, USA. Grasslands such as tallgrass prairies are ideal systems for investigating how seed rain patterns change during community assembly since they are abundant in both early and late-stages of assembly due to restoration efforts (Wilsey, 2021). We revisited the same remnant prairie (Tucker Prairie) as Rabinowitz & Rapp (1980), who conducted the foundational work on seed rain in grasslands. Here, we had the unique opportunity to investigate how seed rain patterns have changed in the last 40+ years. We asked the following questions in our study: Do the timing, quantity, seed mass traits, diversity, and composition of seed rain patterns differ 1) between a remnant prairie measured by Rabinowitz and Rapp (1980) in 1978 versus the same prairie in 2019 and 2) between remnant and restored prairies and change during community assembly? We predicted that seed rain quantity, diversity, and composition in the remnant prairie would be similar across studies, but peaks in dispersal activity would occur earlier due to global climate change advancing plant phenology. As the restoration age increased, we predicted that the timing and quantity of dispersing seeds would become more similar to the remnant prairie since we expected the restorations to establish species dominant in the seed rain. We also predicted the total richness and quantity of dispersing seeds to be greatest in newly restored prairies and decline over time as perennial seeded vegetation displaces introduced and ruderal species from older prairies. Due to tradeoffs between seed rain density and seed mass, we expected average seed mass to also increase with as prairies aged. Lastly, we anticipated that native species diversity and overall species composition would remain divergent from the remnant prairie regardless of restoration age since strongly seed-limited species would not establish well in the prairie restorations.

## Materials and Methods

### Study Sites

Our study sites were at Tucker Prairie (38^○^56’53.6” N, 91^○^59’40.0” W, Callaway County, MO) and at Prairie Fork Conservation Area (38^○^58’29.7” N, 91^○^44’03.3” W, Callaway County, MO)(Fig. S1 A). Tucker Prairie is a 59-hectare tract of unplowed North American tallgrass claypan prairie. Less than 0.5 % of intact tallgrass prairie ecosystem (i.e., never-been-plowed) remains in Missouri, and Tucker Prairie represents the last sizable claypan remnant prairie in north central Missouri (Samson & Knopf, 1994). More than 250 species of plants inhabit Tucker Prairie, with representatives from 57 families and over 150 genera (*Tropicos*, 2023). From 1958 to 2002, Tucker Prairie was burned once every four years in the late winter or early spring (Rabinowitz & Rapp, 1980). Since 2002, Tucker Prairie has been managed on a 5-year burn rotation, where units are burned once in the late winter to early spring (Jan. – Mar.) and again 2-3 years later in the late summer to early fall (Aug. – Oct.). Tucker Prairie was burned one year prior to our study and three years prior to Rabinowitz & Rapp (1980).

Prairie Fork Conservation Area (PFCA) is over 450 hectares of former agricultural land being restored to tallgrass prairie and savanna ecosystems (Newbold et al., 2019). From 2004 onwards, 16-25 hectares are newly seeded each year, with approximately 179 native prairie species collected from Tucker Prairie and other nearby remnant prairies (13.4 to 18.2 kg/ha) (Newbold et al., 2019). As a result, PFCA possesses a chronosequence of reconstructed tallgrass prairies comprised of Tucker Prairie descendants. Reconstructions are managed using a 2-4-year burn schedule (Newbold et al., 2019). For additional details on PFCA management, see Newbold et al. 2019. To capture changes in seed rain dynamics during the restoration process, we grouped our reconstructed sites into three categories, as defined by Newbold et al. (2019) as being representative of restored prairies at various stages of assembly. We measured the seed rain in an old reconstruction (seeded in 2004; burned in 2017), middle-aged reconstruction (seeded in 2013 and 2014; burned in 2017), and a young reconstruction (seeded in 2017; burned in 2018), which were all prepared using the crop method. Similar to other grassland seed rain studies (e.g., Rabinowitz & Rapp, 1980; Schott & Hamburg, 1997), the effort required to sample seed rain at a sufficient temporal and spatial resolution and the lack of additional comparable study sites limited us to using one site per age class.

### Experimental Setup & Data Collection

In May 2019, we deployed artificial turf grass traps (0.01 m^−2^) in Tucker Prairie and in each of the focal PFCA restorations (four total sites). We used artificial turfgrass traps instead of sticky traps because of their durability and resistance to freezing, an issue encountered by Rabinowitz & Rapp (1980) (Molau & Per Mølgaard, 1996). Dispersing seeds become entangled in the blades of artificial grass and are retained until collection. At each site, we randomly established ten, 5 m long transects (Fig. S1 A). We placed transects ∼50 m apart to reduce spatial autocorrelation since most prairie species have mean predicted dispersal distances < 10 m (Sullivan et al., 2018). We then placed traps at 1 m intervals along each transect and affixed them to the ground using ground staples (five traps per transect, 50 traps per site) (Fig. S1 B). We collected and replaced seed traps every 2 weeks from May 31^st^ to December 12^th^, 2019. All traps for a transect were lumped together at each collection period. After collection, we identified seeds to the lowest taxonomic level possible using identification guides and seed reference collections similar to Rabinowitz & Rapp (1980) (Coons et al., 2019.; Martin & Barkley, 1961; details in Supporting Information). Due to identification level differences between our study and Rabinowitz & Rapp (1980), we elevated morphospecies to the same taxonomic level when comparing richness and composition between studies (e.g., *Carex bushii to Carex sp.*). Furthermore, because we sampled a slightly larger area of seed rain than Rabinowitz & Rapp (1980) (0.5 m^2^ in 2019 vs. 0.32 m^2^ per prairie in 1978), we compared seed rain density instead of number of seeds captured between studies. We obtained seed mass data from the Seed Information Database (SER et al., 2023) and the following sources: Barak et al. (2018), Sullivan et al. (2018), Turner & Rabinowitz (1983), and Zirbel et al. (2017). We weighed seeds for taxa lacking published data (details in Supporting Information).

### Data Analysis

#### Quantity and seed mass traits of seed rain

We fit a generalized linear model (GLM) predicting the total number of seeds for the whole year as a function of site (remnant, young reconstruction, middle-aged reconstruction, and old reconstruction) to detect differences in seed rain quantity between prairies. We used a negative binomial distribution (log link) (“Mass” package; Venables & Ripley, 2002) to account for overdispersion in the data. We then conducted a post hoc analysis using Dunnett-style contrasts (“emmeans” package; Lenth et al., 2023) with a multiple comparison adjustment for three tests to determine whether reconstructed sites significantly differed in the number of seeds dispersing compared to the remnant prairie. Similarly, we compared seed biomass (mg) between sites with a GLM using a gamma distribution (log link) followed by a Dunnett-style contrasts post hoc analysis.

We also calculated the community weight mean (CWM) for seed mass (mg) at each site to determine whether seed mass traits changed during assembly. We accomplished this by adjusting seed mass for every species by weighting the number of seeds captured per species at each transect. Then, we used those values to calculate CWMs representative of the “typical” seed mass encountered at transects. Afterward, we fit a linear model predicting the CWM of seed mass (mg) as a function of site. We omitted all unidentified taxa from analyses involving seed mass (mg).

#### Diversity and composition of seed rain

To determine whether annual seed rain from remnant and reconstructed prairies differed in richness, we fit a GLM using a Poisson distribution (log link) predicting total richness as a function of site as before. We then conducted another post hoc analysis using Dunnett-style contrasts to compare reconstruction ages to the remnant, correcting for three tests. We used a similar model and post hoc test to analyze differences in native morphospecies richness between remnant and reconstructed prairies. We also fit a linear model for predicting Shannon diversity index as a function of site. We did not use unidentified taxa in our analysis of richness or Shannon diversity index.

To quantify annual seed rain compositional differences between sites, we first created a community distance matrix using Bray-Curtis dissimilarity, which considers standardized changes in species abundance between sites. Before calculating Bray-Curtis dissimilarity, we relativized species abundance by site total (across rows). We then visualized compositional differences with non-metric multidimensional scaling (NMDS) ordination using two dimensions. We used the envfit function (permutations = 5000) in the “vegan” package (Oksanen et al., 2020) to identify species associated with the compositional differences among sites and plotted them as vectors (p < 0.001).

Using permutational multivariate analysis of variance (PERMANOVA, permutations = 999) and post hoc pairwise comparison tests (“pairwiseAdonis” package; Martinez Arbizu, 2020), we examined potential differences in species composition between remnant and reconstructed prairies. To account for multiple comparisons, we adjusted p-values using a Bonferroni correction. We used the “vegan” package (Oksanen et al., 2020) to conduct all multivariate analyses. We conducted all analyses and visualizations using R (version 4.2.2) and RStudio (version 2023.06.1+524).

## Results

### Quantity, seed mass traits, and timing of seed rain

We captured 121,163 seeds representing 129 morphospecies and 34 plant families across all prairies sampled between May 31st and December 12th, 2019. Estimated seed rain density for every prairie in 2019 far exceeded the estimate reported by Rabinowitz and Rapp (1980) for a remnant tallgrass prairie in 1978 (Table 1). At the same remnant prairie, we captured over two times the density of seed rain in 2019 than in 1978. The quantity and biomass of seed rain in reconstructions decreased with age (Fig. 1A, B). By at least 5-6 years post-initial seeding, the number and biomass of seeds dispersing in reconstructions was comparable to the remnant prairie in 2019. Only the youngest reconstruction had significantly more seeds and biomass dispersing than the remnant prairie sampled in the same year (Fig. 1 A, B; Table S1). We found no significant change in CWM seed mass between sites (Fig. 1 C, F_3,36_ = 1.29, R^2^ = 0.097, p = 0.29).

**Figure 1.**
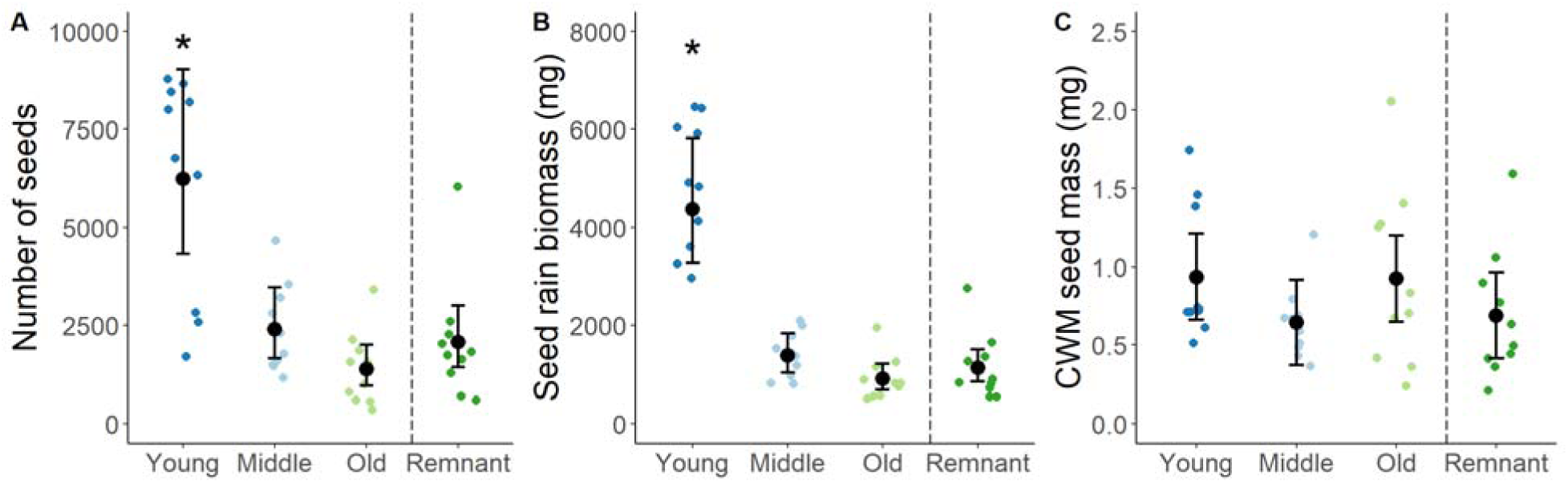
**A)** The number of seeds, **B)** seed rain biomass (mg), and **C)** CWM seed mass (mg) captured per transect in the remnant prairie in 2019 (dark green), young (2-year-old, dark blue), middle-aged (5-6-year-old, light blue), and old (15-year-old, light green) reconstructed prairies. Error bars represent 95% confidence intervals around model estimates (black). Asterisks indicate sites with significant differences from the remnant prairie (p < 0.05).

**Table 1.** Attributes of seed rain captured in a young (2-year-old), middle-aged (5-6-year-old), and old (15-year-old) prairie reconstruction and the same remnant prairie in 1978 and 2019. Characteristics of seed rain in 1978 were obtained from Rabinowitz and Rapp (1980). Numbers outside parentheses indicate the number of morphospecies we successfully identified. Numbers inside parentheses reflect the number of morphospecies observed using the same taxonomic level of identification as Rabinowitz and Rapp (1980).

| Site | Total seeds | Seed rain density<br>(Seeds m<sup>-2</sup> year<sup>-1</sup>) | Total<br>morphospecies<br>richness | Seeds identified<br>(%) |
| --- | --- | --- | --- | --- |
| **Young** | 62,403 | 124,806 | 87 (83) | 99.99 |
| **Middle** | 24,077 | 48,154 | 82 (77) | 99.95 |
| **Old** | 13,876 | 27,752 | 68 (64) | 99.97 |
| **Remnant 2019** | 20,807 | 41,614 | 76 (72) | 99.94 |
| **Remnant 1978 \*** | 6,597 | 20,740 | -- (32) | 99.51 |

\*Trapped seeds in 9 cm diameter sticky traps (n = 50) covered in tanglefoot (Rabinowitz & Rapp, 1980). Reported arithmetic mean for seed rain density.

The timing of seed rain in all prairies exhibited a bimodal pattern of dispersal, where seed rain density peaked once in the early summer and again in the fall (Fig. 2). However, compared to 1978, the timing of peak dispersal activity occurred earlier in 2019. The timing of the seed rain converged with the remnant prairie (2019) as the prairie reconstructions matured. As expected, the youngest reconstructed prairie was the most divergent, where the timing of the first peak in seed rain was delayed by a month compared to the other prairies sampled in 2019. Overall, the timing of the seed rain captured in the middle-aged and old reconstructed prairie closely resembled the timing of the remnant prairie in 2019.

**Figure 2.**
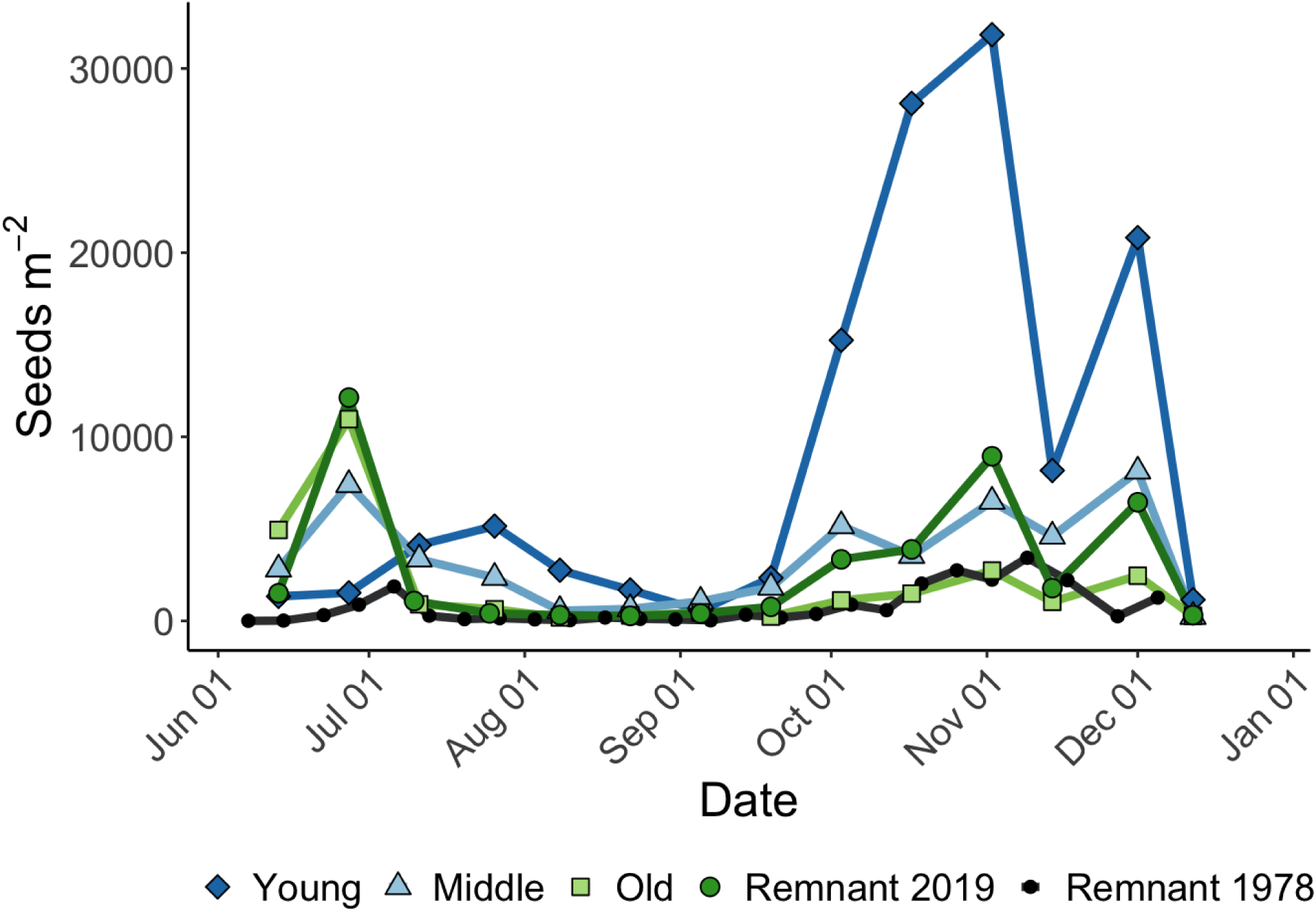
Timing of seed rain in reconstructed prairies of various ages and a remnant prairie. Data for the remnant in 1978 was obtained from Rabinowitz and Rapp (1980).

### Diversity and composition of seed rain

Mean total richness was significantly greater in the young and middle-aged reconstructed prairie compared to the remnant (2019) (Fig. 3 A, Table S1). However, all reconstructed prairies had comparable native morphospecies richness to the remnant prairie in 2019 (Fig. 3 B, Table S1). Total and native richness also trended upward during the study period, with more morphospecies captured in the fall than in the summer (Fig. 3 C, D). All prairies sampled in 2019 had far greater morphospecies captured than the remnant prairie in 1978 (Table 1). In fact, the remnant in 2019 had over twice the number of morphospecies present. All sites shared similar mean Shannon diversity index values ranging from 1.81 ± 0.44 in the youngest reconstruction to 2.28 ± 0.28 in the middle-aged reconstruction (F_3,36_ = 2.637, R^2^ = 0.18, p = 0.064).

**Figure 3.**
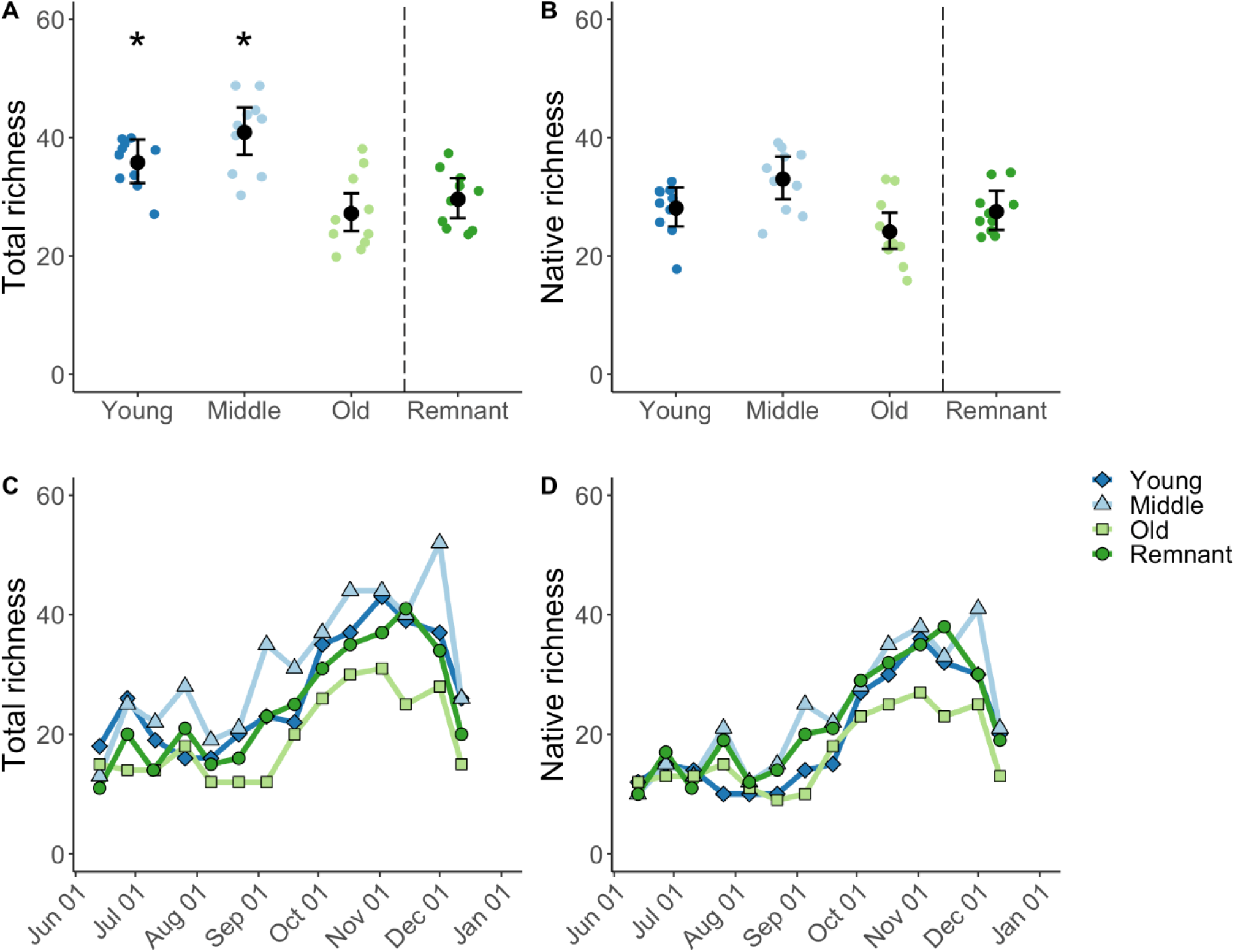
The total (A, C) and native (B, D) morphospecies richness of seed rain captured per transect in the remnant prairie in 2019 (dark green) and the young (2-year-old, dark blue), middle-aged (5-6-year-old, light blue), and old (15-year-old, light green) reconstructed prairies. Error bars represent 95% confidence intervals around model estimates (black). Asterisks indicate sites with significant differences from the remnant prairie (p < 0.05).

Species composition of seed rain significantly differed between all sites (Fig. 4, Table S2). Reconstructed prairies of comparable ages shared the most similarities regarding species composition. Except for some marginal overlap with the oldest reconstruction, the remnant prairie contained a distinct community of dispersing seeds compared to reconstructed sites. Based on the species vectors (Fig. 4), life history played an important role in compositional changes across the reconstructed prairies. Influential annual/biannual species in the seed rain aligned with younger reconstructions and perennial species with older reconstructions and the remnant prairie.

**Figure 4.**
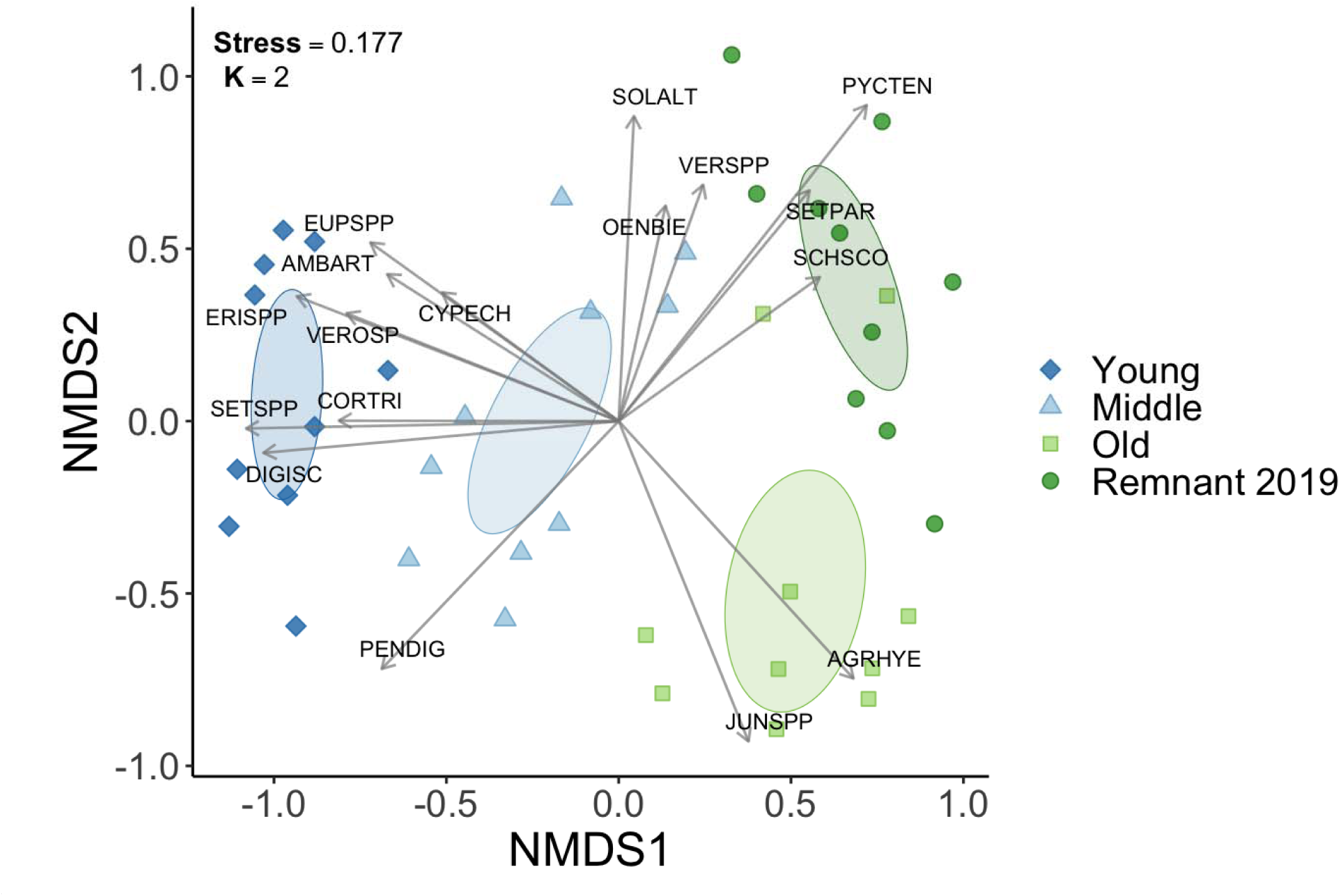
Non-metric multidimensional scaling analysis (NMDS) ordination using Bray-Curtis dissimilarity visualizing seed rain species composition at a remnant prairie in 2019 and prairie reconstructions of various ages. Ellipses represent 95% confidence intervals calculated from standard error. Plot vectors represent species significantly associated with seed rain composition (p ≤ 0.001, see Table S4 for names).

Most of the seeds captured at each site consisted of a handful of dominant species, with many rare species contributing few seeds (Fig. S2). For example, the top three most frequently captured species represented 63.1% of the total seed rain for the remnant prairie in 2019, 76.9% for the remnant prairie in 1978, 63.9% for the old reconstruction, 42.0% for the middle-aged reconstruction, and 63.7% for the young reconstruction. Dominant species in the youngest reconstruction were fecund, disturbance-tolerant species, including smooth crabgrass *Digitaria ischaemum* and boneset *Eupatorium sp.* However, 15 years post-initial seeding, many of these species were almost absent in the oldest reconstruction and similarly rare in the remnant prairie. Instead, native perennial graminoids (e.g., rushes *Juncus sp.*, yellow prairie grass *Sorghastrum nutans*) and forbs (e.g., mountain mint *Pycnanthemum tenuifolium*, foxglove beardtongue *Penstemon digitalis*) dominated the seed rain in the older reconstructions and the remnant prairie (2019 and 1978). Dominant species in the seed rain did not necessarily reflect the dominant species in the vegetation. For example, rushes Juncus sp. were abundant in the seed rain of older prairies but rare in the vegetation (see Supplemental Information, Fig. S4).

## Discussion

Our work demonstrated that seed rain patterns are dynamic in grassland ecosystems and change during community assembly. We report a record amount of seed rain in a grassland ecosystem, where we caught 124,806 seeds m^−2^ year ^−1^ (97.24 g m^−2^ year ^−1^) in the 2-year-old prairie, a number far exceeding most seeding densities of tallgrass prairie restorations, grassland seed addition experiments, and previously published grassland seed rain studies (Myers & Harms, 2009; Rowe, 2010; Rabinowitz & Rapp, 1980; Schott & Hamburg, 1997; Willand et al., 2015; West & Durham, 1996). Despite observing declines in seed rain quantity and biomass over the chronosequence, we did not observe a corresponding increase in CWM seed mass, suggesting that small and large-seeded species are arriving at the same microsites in Missouri prairies of all ages. Although the timing, quantity, and diversity of seed rain in older restored prairies were comparable to the remnant prairie, we found seed rain composition of restored prairies remained divergent from the remnant community even 15 years after planting. Each prairie contained a distinct community of dispersing species despite being initially seeded with species collected from the remnant prairie. Because seed limitation is a common occurrence in grasslands, species missing from the seed rain will likely remain absent due to insufficient dispersal from source populations (Clark et al., 2007; Myers & Harms, 2009; Turnbull et al., 2000).

### Differences in seed rain at the remnant prairie

We found three major differences in seed rain patterns at the same remnant prairie used by Rabinowitz & Rapp (1980). First, peak dispersal activity shifted to occur earlier over the 41 years between studies. Second, the abundance and diversity of seed rain considerably increased since 1978. Lastly, we observed major shifts in seed rain composition between 1978 and 2019. Overall, these findings provide evidence that seed rain patterns are highly variable even within the same mature community.

As expected, we observed an advancement in dispersal phenology for both peaks in dispersal activity in 2019 compared to 1978, where the first and second peaks occurred two weeks and one week earlier, respectively. Plant phenology is sensitive to global climate change, particularly for early-season species (Sherry et al., 2007; Zettlemoyer et al., 2021). Because timing of dispersal activity in conjecture with climatic conditions is likely pivotal for plant recruitment and coexistence outcomes, changes in dispersal phenology could have widespread consequences (DiVittorio et al., 2007; Forrest & Miller-Rushing, 2010). Additional community-level seed rain studies will help reveal whether our findings represent typical interannual variation in peak dispersal activity or long-lasting phenological change.

Secondly, our work revealed significant differences in seed rain quantity and diversity at the same remnant prairie 41 years after Rabinowitz & Rapp’s (1980) foundational study, despite sampling the same number of slightly larger seed traps (100 cm^2^ – 2019, 63.6 cm^2^ – 1978). Against our initial predictions, we captured twice the amount of seed input and morphospecies in 2019 than in 1978. Increases in seed rain quantity and richness between studies were not a result of corresponding long-term increases in aboveground flora richness (see Supplemental Information), but could reflect a change in overall species composition (Fig. S4). Additionally, differences in prescribed fire timing (burned 1 year prior to 2019, 3 years prior to 1978) and precipitation during months critical for seed production (above average in 2019, below average in 1978) presumably contributed to the substantial changes in seed rain density and richness observed between studies since these factors can stimulate the flowering and reproductive output of prairie species (Table S3) (Daubenmire, 1968; Rabinowitz et al. 1989; Wagenius et al. 2020). Due to the large interannual variation observed between studies, we encourage future work to explore how climatic conditions and fire influence community-level seed production.

Lastly, we observed shifts in the dominant species composition of seed rain captured between 1978 and 2019, suggesting major compositional changes have occurred between studies (Rabinowitz & Rapp, 1980) (Fig. S3). For example, mountain mint *Pycnanthemum tenuifolium*, a species rare in the aboveground flora and absent from the seed rain in 1978 (Rabinowitz & Rapp, 1980; Rabinowitz et al. 1981), was the most frequently captured species in 2019 at the same prairie. Despite catching twice the number of morphospecies, we were still unable to recapture 28.1% of the species caught in the 1978 study (Rabinowitz & Rapp, 1980). Because we used artificial turfgrass traps instead of sticky traps, trap design could have contributed to the observed differences in seed rain patterns between years. However, concurrent changes in the aboveground flora over a 38-year period at the remnant prairie provides additional support that differences in seed rain composition between studies does not represent typical interannual variation or methodology differences but rather long-term change (see Supplemental Information, Fig. S4). Since species establishment and richness are dependent on intra- and interspecific propagule supply, changes in annual seed input can have major implications for the future community composition of the aboveground flora (DiVittorio et al., 2007; Myers & Harms, 2009).

Many studies have cited Rabinowitz & Rapp’s (1980) findings as a high estimate for seed rain, not only in grasslands but for all ecosystems (e.g., Jochems et al., 2022; Kettenring & Galatowitsch, 2011; Martin & Wilsey, 2006; Myers & Harms, 2009). Although it is unclear whether 2019 represents a typical or extreme year, our results show that grasslands can produce far more abundant and diverse seed rain than previously thought. Seed production in remnant prairies is likely even greater than our findings suggest due to trap obstruction from vegetation (Brown & Cahill Jr., 2020), post-dispersal seed predation (Johnson & Zettlemoyer, 2022), and secondary dispersion of seeds by wind, water, and animals (Chambers & MacMahon, 1994). Because our trapping design did not exclude animals, animal behavior may have influenced species representation in our estimates. For example, species favored by animal predators (e.g., prairie dropseed *Sporobolus heterolepis*) or reliant on animal dispersers (e.g., wild strawberry *Fragaria virginica*) were likely underrepresented in our traps as evidence by their high presence in the aboveground flora and low to no presence in our seed rain samples (Figure S4) (Parker-Smith, 2022). Regardless, we captured over 20 times the amount of seed biomass, including unassisted, wind, and animal-dispersed species, in the remnant in 2019 (22.92 g m^−2^) compared to typical restoration seeding rates (Rowe, 2010). Long-lasting differences from remnant aboveground vegetational communities in restored prairies may result from continued seed limitation.

### Differences in seed rain across a restoration chronosequence

We found considerable differences in seed rain patterns across a time-since-restoration gradient of tallgrass prairie restorations. Seed rain captured in the newly restored prairie was significantly more abundant and diverse compared to older restorations; however, increases in richness were solely due to introduced species. Each restored prairie along the chronosequence had a distinct community of dispersing seeds with notable changes in dominant species over time. Consequently, the timing of dispersal activity reflected the phenology of dominant species, which differed most in the youngest prairie restoration compared to the older restorations. Our results indicate that key aspects of the seed rain change substantially as restored communities age.

Observed changes in seed rain diversity and composition across a restoration gradient closely mirrored well-established patterns in the vegetative community, where prairie restorations become increasingly dominated by perennial grasses over time. Seed rain in young prairie restorations was dominated by introduced, annual, and disturbance-tolerant species that were eventually displaced by native perennial species 15 years after restoration efforts began, resulting in the oldest restoration losing total richness. Unlike other studies that saw a reduction in native richness in the aboveground plant community of older restorations (e.g., Hansen & Gibson, 2014; Sluis, 2002), we captured comparable native richness of dispersing seeds in all restored prairies. However, the identities of these native species changed as the prairies aged. Overall, a combination of native species turnover and displacement of introduced species resulted in distinct seed rain compositions across the chronosequence.

Although many studies observe a negative relationship between seed rain density and seed size at the species level (Moles et al., 2004; Moles & Westoby, 2006; Turnbull et al., 1999), we found that this relationship does not extend to the community level. Against our expectations, small and large-seeded species were similarly present in the seed rain of restored communities of all ages despite the incredible quantity of seeds dispersing in early-successional prairies, indicating a lack of a numeric competitive advantage to small-seeded species. Instead, changes in seed rain quantity occurred because of compositional shifts toward long-lived perennial species. Given the well-documented evidence that large-seeded species tend to be seed limited (Clark et al., 2007), our results support the role of post-dispersal processes that limit recruitment of large-seeded species (e.g., consumers, disease, etc.) (Chambers & MacMahon, 1994). Additionally, introduced species were also prevalent in the younger restorations, which are often more fecund than native congeners and may explain differences in seed input across the chronosequence of restorations (Pyšek & Richardson, 2007). Other grassland seed rain studies similarly report denser seed input in young communities compared to older restored or remnant communities (Kettenring & Galatowitsch, 2011; West & Durham, 1991), suggesting that larger quantities of seed input may be characteristic of early-assembling grassland communities.

### Implications for Research and Management

We demonstrated that tallgrass prairie seed rain is far more diverse and abundant than previously estimated. Furthermore, we showed that seed dispersal patterns change during community assembly, with young communities having significantly greater propagule pressure than older communities. Together, our research supports that restoration efforts have yet to fully replicate the community composition seen at remnant prairies (Barak et al., 2017; Newbold et al., 2019; Sluis et al., 2018). Species only present in the remnant prairie seed rain were often understory species like yellow star grass *Hypoxis hirsuta* and arrow-leaf violet *Viola sagittata*, which are difficult to establish and highly desirable in restorations (Barak et al., 2022). Because these species rely on seed dispersal to arrive in new communities, they will likely remain absent without human assistance (Sperry et al., 2019). Restoration efforts aim to overcome seed limitation by broadcast seeding at rates based on studies of remnant prairie seed rain such as Rabinowitz & Rapp (1980) (Rowe, 2010). However, we captured over twice the amount of seed input in the same remnant and over six times the amount in the newly restored prairie as Rabinowitz & Rapp (1980). New species establishment from seed dispersal is an inherently probabilistic process, where increasing propagule supply correspondingly increases the chances of successfully entering the community (D’Antonio et al., 2001; DiVittorio et al., 2007). Therefore, increasing restoration seeding rates may be needed to meet restoration composition and diversity goals since current rates may not compete with the propagule pressure of undesirable species found in newly restored prairies.

## Author Contributions

L.L.S and K.C.W conceived the ideas and methodology. K.C.W, L.L.S, M.J.P., and E.M.E. all contributed towards data collection. K.C.W analyzed the data. K.C.W. led the writing of the manuscript, and all authors contributed to manuscript formation and editing.

## Acknowledgements

We thank Prairie Fork Conservation Area for allowing us to use their property and Chris Newbold, Jeff Demand, Amber Edwards, and Melody Kroll for their help with site selection and information. We thank Larissa Kahan, Savana Presson, Kelsey Jaeger, Danielle Gafford, and Christian Perez-Martinez for assistance with data collection, John Snyder for providing valuable data analysis input, and Deborah Finke, Elizabeth King, Manuel Leal, Lars Brudvig, Gaurav Kandlikar, the Sullivan Lab, Pieter De Frenne, and two anonymous reviewers for providing excellent feedback on our manuscript. We also thank James Carrel for graciously providing us with floristic survey data of Tucker Prairie in 1981. The Prairie Fork Charitable Endowment Trust and Long-Term Agroecosystem Research (LTAR) network (58-5070-9-016 and 58-5070-2-018) provided funding that supported our research. LTAR is supported by the United States Department of Agriculture. This is KBS contribution #2386.

