## supplemental information for "Quantifying seed rain patterns in a remnant and a chronosequence of restored tallgrass prairies in north central Missouri"

**Supplemental Information on Methods**

*Processing of seed traps*

 After collection, all traps from a transect and sampling period were grouped together and sieved through a series of soil sieves (1 mm, 500 μm, 250 μm mesh). We then counted and identified captured seeds in sieved layers. Because of their extremely small size, we estimated the number of rush seeds (*Juncus sp.*) when there were over 200 seeds present in a sample. We first sieved the samples through 180 μm and 150 μm mesh soil sieves. Then we subsampled each sieved layer by calculating the average number of rush seeds per 1 cm^2^ area (n = 3) and multiplying that average by the total area covered by the sample layer. Lastly, we summed the number of estimated rush seeds per layer to calculate the total number of rush seeds in a sample.

Seed Mass Data

We also individually weighed seeds (21 per species) to obtain the mean one seed mass (mg) for taxa lacking accessible data. We were unable to obtain seed mass for two taxa because of their rarity and lack of published data in the literature. For species with multiple reported masses, we converted all weights to one seed mass (mg) and used the average value for analysis. When we could only identify taxa to genus level, we calculated the average seed mass for all members of that genus known to inhabit our study sites (Newbold et al., 2019; *Tropicos*, 2023).

*Comparison of vegetation in the remnant prairie between 1981 and 2019*

To determine whether the remnant prairie also experienced long-term changes in aboveground flora diversity and composition, we compared floristic survey data collected in 1981 to data we collected in 2019. In 1981 (Jul. – Aug.), the aboveground flora was assessed by counting the number of stems per species in 1 m^2^ frame quadrats regularly spaced at 50 to 150 m intervals across the entire prairie (similar to Drew, 1947). For our analysis, we only used data collected in 1981 from the northern portion of the remnant prairie where our seed rain transects were located, representing a sampling area of 23 m^2^ (n = 23 quadrats). In 2019 (Aug. – Sep.), we sampled the vegetational community in a 1 m^2^ area around each seed trap at transects. Since one transect was dropped from the analysis due to incomplete data, our sampling area represents 45 m^2^ of vegetation (n = 45 quadrats). Because we measured the percent aerial cover of all vascular species rooted within sampling areas instead of the number of stems in 2019, we standardized all floristic data by presence/absence. Additionally, we elevated certain taxa to genus level due to differences in taxonomic identification level between surveys (e.g., *Carex sp*., *Jucus sp.*, etc.). Unidentified taxa were removed from all further analyses.

We assessed whether there were differences in aboveground flora morphospecies richness between 1981 and 2019 by predicting richness as a function of the survey year using a generalized linear model with a Poisson distribution followed by an analysis of deviance test. Furthermore, we visualized long-term changes in species composition using non-metric multidimensional scaling (NMDS) ordination (k = 3, stress = 0.16) followed by a permutational multivariate analysis of variance (PERMANOVA) test using Jaccard distance. We also identified influential species using the envfit() function (permutations = 5000) in the “vegan” package (Oksanen et al., 2020) and plotted them as vectors in our NMDS ordination (p < 0.0002).

**Supplemental Information on Results**

*Comparison of vegetation in the remnant prairie between 1981 and 2019*

We found that the average number of morphospecies did not differ between 1981 (18.9 ± 3.29 species per m^2^) and 2019 (18.2 ± 2.96 species per m^2^) in the same remnant prairie flora (X^2^_1_ = 0.34, p = 0.56). However, the species composition of the aboveground flora changed in the 38 years between surveys (pseudo-F = 12.57, R^2^ = 0.16, p < 0.001).

**Supplemental Figures & Tables**

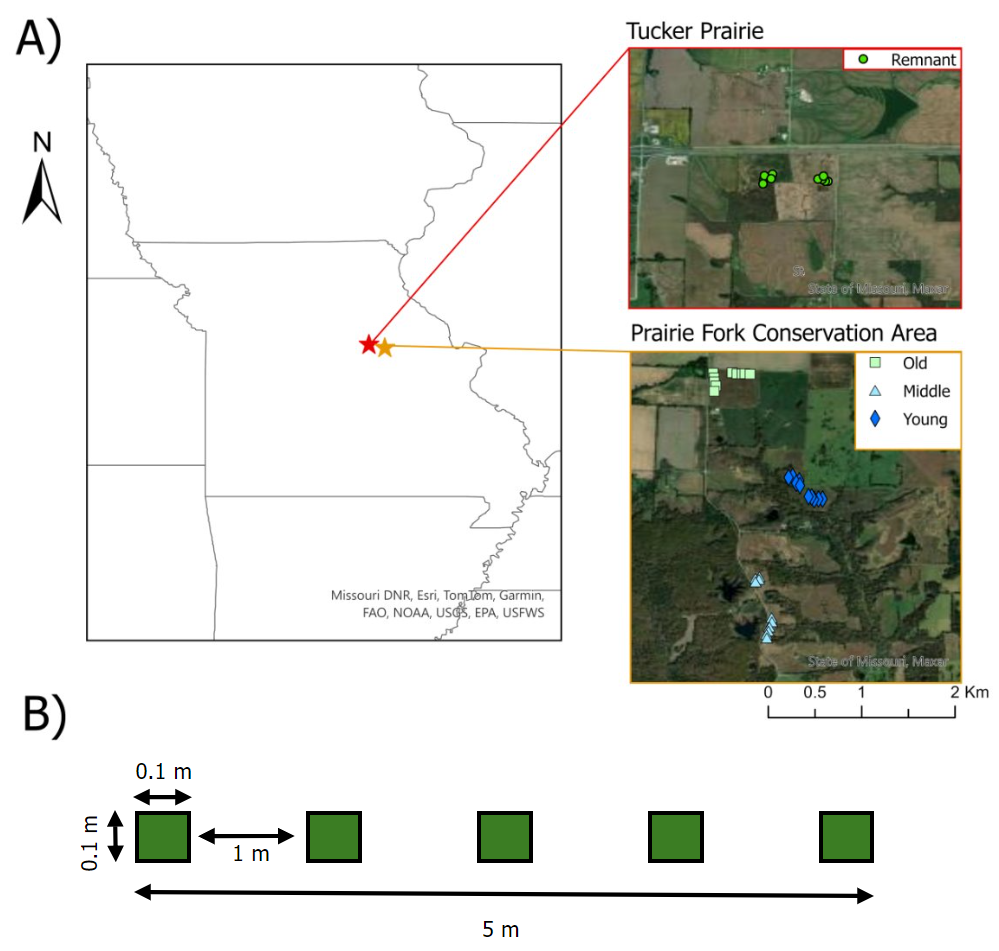

**Figure S1. A)** Map of study sites, Tucker Prairie and Prairie Fork Conservation Area, and transect locations in the old, middle-aged, and young restored and the remnant prairie (n = 10 transects per site; 40 transects total). **B)** Experimental design of transects used to sample seed rain in prairies. Green squares represent artificial turfgrass seed trap locations within a transect (n = 5 traps per transect; 50 traps per site; 200 traps total). Traps were replaced every 2 weeks from May to Dec. 2019.

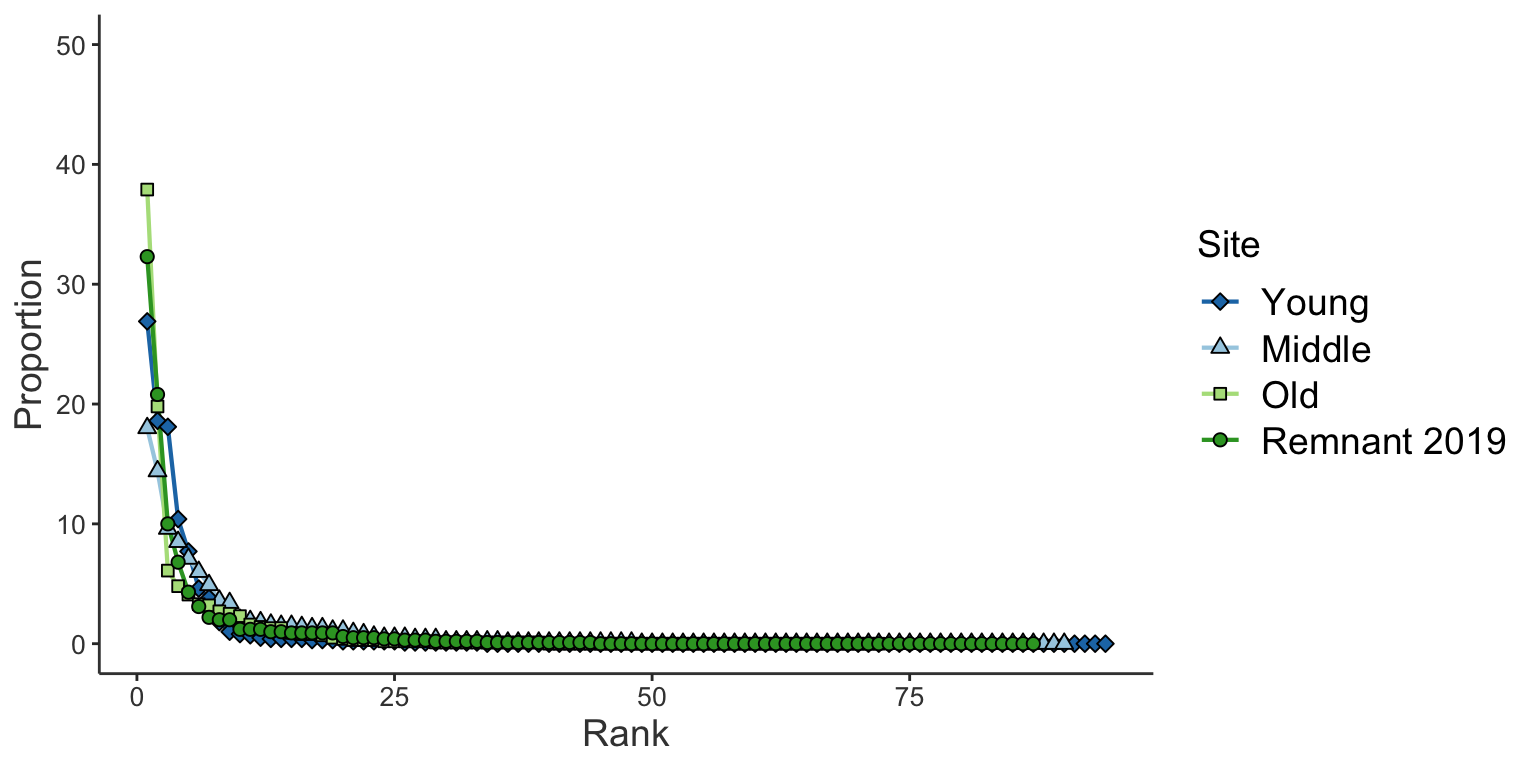

**Figure S2.** Rank abundance curves for a chronosequence of prairie restorations (young = 2-year-old, middle = 5-6-year, and old = 15-year-old) and a remnant prairie in 2019.

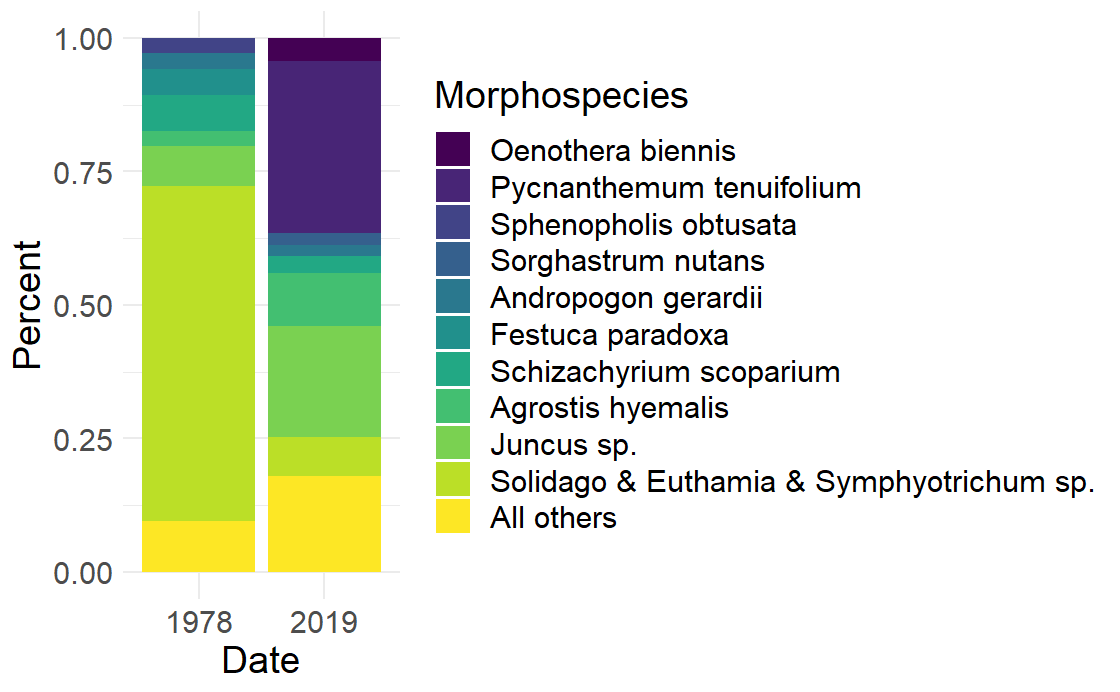

**Figure S3.** Comparison of dominant morphospecies in the seed rain between 1978 (Rabinowitz & Rapp 1980) and 2019 at the same remnant tallgrass prairie. Species that individually contributed less than 2% of the total seed rain during a sampling year were grouped together in the “all others” category.

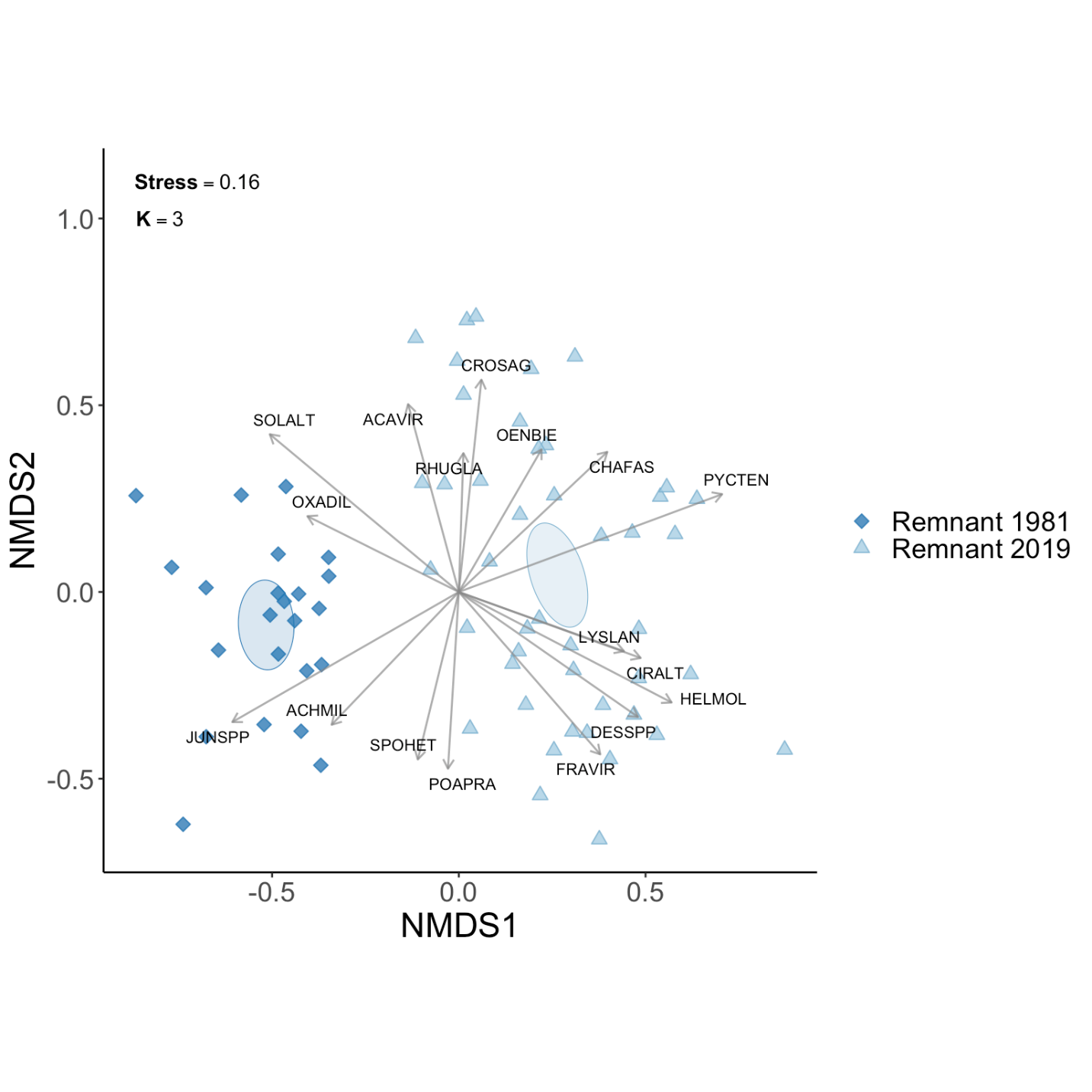

**Figure S4.** Non-metric multidimensional scaling (NMDS) ordination visualizing the vegetational composition of the remnant prairie in 1981 (dark blue diamonds) and 2019 (light blue triangles) in three dimensions. Ellipses represent 95% confidence intervals calculated from standard error. Plot vectors represent species significantly associated with the composition of the aboveground flora (p < 0.0002): ACAVIR = *Acalypha virginica*, ACHMIL = *Achillea millefolium*, CHAFAS = *Chamaecrista fasciculata*, CIRALT = *Cirsium altissimum*, CROSAG = *Crotalaria sagittalis*, DESSPP = *Desmodium sp*., FRAVIR = *Fragaria virginiana*, HELMOL = *Helianthus mollis*, JUNSPP = *Juncus sp*., LYSLAN = *Lysimachia lanceolata*, OENBIE = *Oenothera biennis*, OXADIL = *Oxalis dillenii*, POAPRA = *Poa pratensis*, PYCTEN = *Pycnanthemum tenuifolium*, RHUGLA = *Rhus glabra*, SOLALT = *Solidago altissima*, SPOHET = *Sporobolus heterolepis*.

**Table S1.** Summary results for pairwise contrast tests comparing the mean number of seeds (Z, (P)), mean seed biomass (mg) (T,(P)), mean total morphospecies richness (Z, (P)), and mean native morphospecies richness (Z, (P)) between reconstructed prairies of various ages and the remnant prairie in 2019. Bolded values indicate significance at the α 0.05 level.

| **Variable** | **Young – Remnant** | **Middle – Remnant** | **Old – Remnant** |
| --- | --- | --- | --- |
| Number of seeds | 4.13 **(< 0.001)** | 0.55 (0.548) | -1.52 (0.300) |
| Seed biomass (mg) | **7.48 (<0.001)** | 1.054 (0.58) | -1.06 (0.58) |
| Total richness | 2.46 **(< 0.05)** | 4.28 **(< 0.001)** | - 0.97 (0.634) |
| Native richness | 0.30 (0.964) | 2.28 (0.062) | - 1.45 (0.335) |

**Table S2.** PERMANOVA and pairwise comparison results from comparing seed rain composition between reconstructed prairies of various ages and the remnant prairie in 2019.

| **Source** | **Df** | **SS** | **R^2^** | **F (P)** |
| --- | --- | --- | --- | --- |
| Site | 3 | 5.16 | 0.41 | 8.22 **(< 0.001)** |
| Residual | 36 | 7.54 | 0.59 | - |
| *Pairwise comparison* | | | | |
| Young – Middle | 1 | 1.19 | 0.25 | 6.06 **(< 0.001)** |
| Young – Old | 1 | 2.43 | 0.39 | 11.34 **(< 0.001)** |
| Young – Remnant | 1 | 2.640 | 0.43 | 13.79 **(< 0.001)** |
| Middle – Old | 1 | 1.21 | 0.23 | 5.32 **(< 0.001)** |
| Middle – Remnant | 1 | 1.71 | 0.32 | 8.39 **(< 0.001)** |
| Old – Remnant | 1 | 1.14 | 0.22 | 5.15 **(< 0.01)** |

**Table S3.** Monthly precipitation (mm) and average air temperature (^o^C) in 1978 and 2019 in Callaway County, MO (station USC00234271). Mean values represent the mean and standard deviation (SD) of these factors during the 41-year period between studies. Dashes represent missing values. Data obtained from the National Centers for Environmental Information.

| **Month** | **Precipitation (mm)** | | | **Average Air Temperature (^o^C)** | | |
| --- | --- | --- | --- | --- | --- | --- |
|  | **1978** | **2019** | **Mean ± SD** | **1978** | **2019** | **Mean ± SD** |
| Jan. | 19.81 | 81.02 | 52.98 ± 39.76 | -6.89 | -1.17 | -1.21 ± 2.81 |
| Feb. | 25.65 | 69.09 | 52.53 ± 36.55 | -5.89 | -0.17 | 0.85 ± 3.30 |
| Mar. | 112.78 | 118.62 | 80.44 ± 42.50 | 2.89 | 4.67 | 6.72 ± 2.39 |
| Apr. | 114.05 | 72.64 | 112.71 ± 60.97 | 14.06 | 13.72 | 12.92 ± 1.73 |
| May | 111.76 | 222.00 | 131.12 ± 62.35 | 17.00 | 18.50 | 18.15 ± 1.70 |
| Jun. | 59.18 | 104.65 | 102.27 ± 65.04 | 23.22 | 22.78 | 23.28 ± 1.17 |
| Jul. | 106.42 | 42.16 | 104.7 ± 67.11 | 26.22 | 26.28 | 25.74 ± 1.39 |
| Aug. | 89.66 | 199.14 | 100.91 ± 62.77 | 24.83 | 24.72 | 24.89 ± 1.69 |
| Sep. | 47.24 | - | 94.75 ± 81.99 | 23.00 | 12.78 | 20.36 ± 1.61 |
| Oct. | 56.64 | 188.21 | 92.85 ± 68.14 | 12.89 | 4.94 | 13.58 ± 1.70 |
| Nov. | 121.15 | 54.61 | 83.95 ± 63.56 | 12.89 | 4.11 | 7.04 ± 2.37 |
| Dec. | - | 29.72 | 57.03 ± 40.25 | 8.00 | 0.94 | 0.98 ± 2.61 |
| Total | 864.36 | 1,341.88 | 972.86 ± 321.83 |  |  |  |

**Table S4.** Total number of seeds captured at all sites. Divide by 200 to get mean number of seeds per turf grass trap. Multiply by 0.5 to convert to seeds per m^2^ (total trapping area 2 m^2^).

| **Scientific Name** | **SPP6** | **Jun. 13th** | **Jun. 27th** | **Jul. 10^th^** | **Jul. 11^th^** | **Jul. 26^th^** | **Aug. 8^th^** | **Aug. 22^nd^** | **Sep. 5^th^** | **Sep. 19^th^** | **Oct. 3^rd^** | **Oct. 17^th^** | **Nov. 2^nd^** | **Nov. 14^th^** | **Dec. 1^st^** | **Dec. 12^th^** | **Total** |
| --- | --- | --- | --- | --- | --- | --- | --- | --- | --- | --- | --- | --- | --- | --- | --- | --- | --- |
| *Agrostis hyemalis var. hyemalis* | AGRHYE | 2010 | 2660 | 73 | 53 | 27 |  | 5 | 5 |  |  | 4 | 2 |  |  |  | **4839** |
| *Sphenopholis obtusata* | SPHOBT | 1470 | 1032 | 10 | 73 | 124 | 7 | 2 | 40 | 8 | 1 | 4 | 12 |  | 45 | 1 | **2829** |
| *Juncus sp.* | JUNSPP | 933 | 10818 | 340 | 781 | 412 | 109 | 66 | 135 |  |  |  |  |  | 98 |  | **13692** |
| *Cerastium sp.* | CERSPP | 158 | 71 |  | 12 | 4 | 6 | 1 | 1 | 10 |  | 1 |  |  |  | 4 | **268** |
| *Coreopsis lanceolata* | CORLAN | 143 | 53 |  | 48 | 42 | 1 | 8 | 28 | 15 | 5 | 7 | 9 |  | 6 |  | **365** |
| *Veronica sp.* | VEROSP | 108 | 77 |  | 16 | 7 | 5 | 6 | 2 | 3 | 3 |  |  |  |  |  | **227** |
| *Poa pratensis* | POAPRA | 105 | 242 | 29 | 302 | 118 | 37 | 26 | 31 | 30 | 2 | 6 | 5 | 2 | 3 | 1 | **939** |
| *Plantago virginica* | PLAVIR | 98 | 99 |  | 46 | 14 | 2 | 4 | 5 |  | 7 | 5 | 9 | 8 | 10 | 2 | **309** |
| *Myosotis verna* | MYOVER | 69 | 58 |  | 11 | 5 | 4 | 1 | 6 | 1 | 1 |  | 3 |  | 1 | 1 | **161** |
| *Silene antirrhina* | SILANT | 65 | 9 |  |  |  |  |  |  |  |  |  |  |  |  |  | **74** |
| *Triodanis perfoliata* | TRIPER | 24 | 22 | 3 | 7 | 13 |  |  |  |  |  |  |  |  |  |  | **69** |
| *Carex bushii* | CARBUS | 17 | 67 | 23 | 28 | 41 | 10 | 13 | 12 | 6 | 1 | 4 | 7 | 3 | 5 | 1 | **238** |
| *Alopecurus carolinianus* | ALOCAR | 17 | 1 |  | 2 |  |  |  |  |  |  |  |  |  |  |  | **20** |
| *Carex festucacea* | CARFES | 14 | 222 | 18 | 85 | 58 | 27 | 19 | 134 | 16 | 14 | 7 | 28 | 4 | 8 | 1 | **655** |
| *Geranium carolinianum* | GERCAR | 13 | 15 |  | 2 |  |  |  |  |  |  |  |  |  |  |  | **30** |
| *Anagallis minima* | ANAMIN | 11 | 15 |  | 67 | 13 | 1 |  | 2 | 9 |  |  |  |  |  |  | **118** |
| *Capsella bursa-pastoris* | CAPBUR | 7 | 12 |  |  | 3 |  |  |  |  |  |  |  |  |  |  | **22** |
| *Oxalis dillenii* | OXADIL | 5 | 4 |  | 1 | 3 | 2 | 3 |  |  | 1 | 3 | 11 | 4 | 1 |  | **38** |
| *Penstemon digitalis* | PENDIG | 4 | 1 |  |  | 35 |  | 6 | 8 | 79 | 1238 | 991 | 4386 | 2426 | 7333 | 128 | **16635** |
| *Tradescantia ohiensis* | TRAOHI | 3 | 66 |  | 33 | 12 |  | 7 |  |  | 1 |  | 1 | 1 |  |  | **124** |
| *Galium aparine* | GALAPA | 3 | 9 |  |  | 1 |  |  |  |  |  |  |  |  |  |  | **13** |
| *Hordeum pusillum* | HORPUS | 3 | 7 |  |  |  |  |  |  |  |  |  |  |  |  |  | **10** |
| *Dichanthelium lanuginosum* | DICLAN | 2 | 68 | 25 | 1 | 14 | 14 | 11 | 11 | 19 | 3 | 9 | 9 | 5 | 8 | 3 | **202** |
| *Lepidium virginicum* | LEPVIR | 2 | 11 |  | 3 | 1 |  |  |  |  |  |  |  |  |  |  | **17** |
| *Unknown spp* | UNKSPP | 2 | 4 | 1 |  | 3 |  | 2 | 10 | 3 | 3 | 1 | 3 | 1 |  | 1 | **34** |
| *Thlaspi arvense* | THLARV | 2 | 4 |  |  |  |  | 1 | 4 |  |  |  | 4 | 2 | 2 | 1 | **20** |
| *Ratibida pinnata* | RATPIN | 2 | 1 |  | 1 | 13 |  | 2 | 36 | 38 | 157 | 120 | 167 | 62 | 136 | 5 | **740** |
| *Eryngium yuccifolium* | ERYYUC | 1 | 1 |  | 3 | 1 | 7 | 1 | 9 | 6 | 51 | 109 | 170 | 50 | 82 | 4 | **495** |
| *Melilotus sp.* | MELSPP | 1 |  |  | 22 | 22 | 51 | 162 | 125 | 156 | 62 | 22 | 16 | 10 | 7 |  | **656** |
| *Hypoxis hirsuta* | HYPHIR | 1 |  |  |  |  |  |  |  |  |  |  |  |  |  |  | **1** |
| *Platanus occidentalis* | PLAOCC | 1 |  |  |  |  |  |  |  |  |  |  |  |  |  |  | **1** |
| *Erigeron sp.* | ERISPP |  | 148 | 8 | 2414 | 2924 | 1383 | 694 | 64 | 87 | 40 |  |  |  |  |  | **7762** |
| *Vulpia octoflora* | VULOCT |  | 34 |  | 7 | 4 | 2 | 2 |  |  | 1 |  | 1 |  |  |  | **51** |
| *Setaria sp.* | SETSPP |  | 30 |  | 11 |  | 1 | 5 | 23 | 268 | 2069 | 1689 | 1094 | 177 | 242 | 25 | **5634** |
| *Digitaria ischaemum* | DIGISC |  | 22 |  | 2 | 1 | 6 | 5 | 14 | 469 | 4734 | 8006 | 3838 | 955 | 592 | 165 | **18809** |
| *Monarda fistulosa subsp. fistulosa* | MONFIS |  | 18 |  |  | 4 | 4 | 8 | 35 | 32 | 109 | 25 | 48 | 18 | 64 |  | **365** |
| *Lobelia spicata* | LOBSPI |  | 16 |  |  | 1 |  |  |  |  |  |  |  |  |  |  | **17** |
| *Achillea millefolium* | ACHMIL |  | 14 |  |  | 15 | 22 | 41 | 24 | 19 | 31 | 28 | 64 | 47 | 34 | 6 | **345** |
| *Coreopsis palmata* | CORPAL |  | 13 |  | 17 | 3 | 1 |  |  |  |  |  |  |  |  |  | **34** |
| *Barbarea vulgaris* | BARVUL |  | 11 | 2 | 2 | 12 | 6 | 6 | 11 | 5 | 12 | 11 | 97 | 11 | 45 | 4 | **235** |
| *Festuca arundinacea* | FESARU |  | 9 |  | 3 |  | 1 |  | 1 |  |  |  |  |  |  |  | **14** |
| *Carex cephalophora* | CARCEP |  | 8 |  |  |  |  |  |  |  |  |  |  |  |  |  | **8** |
| *Medicago lupulina* | MEDLUP |  | 7 |  | 52 | 97 | 65 | 78 | 61 | 35 | 10 | 8 |  | 1 | 5 |  | **419** |
| *Rubus sp.* | RUBSPP |  | 4 | 2 |  |  | 34 |  |  |  |  | 1 | 1 |  | 1 |  | **43** |
| *Bromus japonicus* | BROJAP |  | 4 |  | 5 | 2 | 28 | 5 | 5 | 4 |  |  |  |  | 25 | 1 | **79** |
| *Pycnanthemum tenuifolium* | PYCTEN |  | 3 | 3 |  | 7 | 3 | 5 | 50 | 407 | 1087 | 861 | 2726 | 761 | 2798 | 100 | **8811** |
| *Carex bicknellii* | CARBIC |  | 3 |  |  |  |  |  |  |  |  |  |  |  | 1 |  | **4** |
| *Ambrosia artemisiifolia* | AMBART |  | 2 |  |  | 2 | 1 |  | 4 | 1 | 18 | 26 | 13 | 4 | 8 | 2 | **81** |
| *Vernonia sp.* | VERSPP |  | 2 |  |  |  |  |  |  | 3 | 16 | 39 | 27 | 84 | 17 | 15 | **203** |
| *Rumex crispus* | RUMCRI |  | 2 |  |  |  |  |  |  |  |  |  |  |  |  |  | **2** |
| *Koeleria macrantha* | KOEMAC |  | 1 |  | 65 | 109 | 17 | 10 | 58 | 7 |  |  |  |  | 1 |  | **268** |
| *Kummerowia sp.* | KUMSPP |  | 1 |  |  |  |  | 2 | 4 | 2 | 22 | 55 | 81 | 59 | 130 | 12 | **368** |
| *Verbena hastata* | VERHAS |  | 1 |  |  |  |  |  |  |  |  |  | 4 |  | 5 | 1 | **11** |
| *Cardamine sp.* | CARDSP |  | 1 |  |  |  |  |  |  |  |  |  |  |  |  |  | **1** |
| *Persicaria longiseta* | PERLON |  | 1 |  |  |  |  |  |  |  |  |  |  |  |  |  | **1** |
| *Taraxacum officinale* | TAROFF |  | 1 |  |  |  |  |  |  |  |  |  |  |  |  |  | **1** |
| *Viola sagittata* | VIOSAG |  | 1 |  |  |  |  |  |  |  |  |  |  |  |  |  | **1** |
| *Polygala sp.* | POLSPP |  |  | 3 |  |  | 3 | 2 | 7 | 1 | 1 | 1 |  |  |  |  | **18** |
| *Schizachyrium scoparium* | SCHSCO |  |  | 2 |  | 1 | 2 |  | 2 | 16 | 242 | 274 | 372 | 103 | 127 | 29 | **1170** |
| *Scirpus pendulus* | SCIPEN |  |  |  | 7 | 17 |  |  |  |  |  | 1 | 5 | 1 | 2 |  | **33** |
| *Scleria triglomerata* | SCLTRI |  |  |  | 4 |  |  |  |  |  |  |  |  |  |  |  | **4** |
| *Blephilia ciliata* | BLECIL |  |  |  | 2 |  |  |  |  |  |  |  |  |  |  |  | **2** |
| *Andropogon gerardii* | ANDGER |  |  |  | 1 |  |  |  | 23 | 3 | 63 | 121 | 287 | 31 | 45 | 3 | **577** |
| *Rudbeckia hirta var. pulcherrima* | RUDHIR |  |  |  |  | 36 | 1 |  | 13 | 25 | 193 | 82 | 238 | 82 | 129 | 21 | **820** |
| *Cyperus acuminatus* | CYPACU |  |  |  |  | 20 |  | 8 | 30 | 5 | 12 | 5 | 9 | 4 | 10 |  | **103** |
| *Carex annectens* | CARANN |  |  |  |  | 15 |  |  |  |  |  |  |  |  |  |  | **15** |
| *Festuca paradoxa* | FESPAR |  |  |  |  | 5 | 5 | 9 | 11 | 18 | 161 | 13 | 12 | 4 | 7 | 5 | **250** |
| *Setaria parviflora* | SETPAR |  |  |  |  | 4 | 1 | 8 |  | 4 | 4 | 6 | 6 | 1 | 4 | 2 | **40** |
| *Amorpha canescens* | AMOCAN |  |  |  |  | 4 | 1 | 1 |  | 1 | 1 |  | 1 | 2 | 13 |  | **24** |
| *Cyperus echinatus* | CYPECH |  |  |  |  | 3 | 22 | 181 | 114 | 17 | 14 | 31 | 30 | 23 | 30 | 7 | **472** |
| *Linum sulcatum var. sulcatum* | LINSUL |  |  |  |  | 3 | 7 | 3 |  |  |  | 4 | 16 | 6 | 16 | 4 | **59** |
| *Galium obtusum subsp. obtusum* | GALOBT |  |  |  |  | 3 | 4 | 6 | 9 | 4 |  | 1 |  | 4 | 6 |  | **37** |
| *Lythrum alatum* | LYTALA |  |  |  |  | 2 |  |  |  |  |  |  |  |  |  |  | **2** |
| *Acalypha virginica* | ACAVIR |  |  |  |  |  | 3 | 1 | 10 | 28 | 49 | 25 | 16 |  | 2 |  | **134** |
| *Crotalaria sagittalis* | CROSAG |  |  |  |  |  | 2 | 30 | 28 | 48 | 37 | 24 | 11 | 4 | 1 | 2 | **187** |
| *Dianthus armeria* | DIAARM |  |  |  |  |  | 1 |  |  |  |  |  |  |  | 2 |  | **3** |
| *Mollugo verticillata* | MOLVER |  |  |  |  |  |  | 2 | 1 |  |  |  | 1 |  | 8 |  | **12** |
| *Helianthus mollis* | HELMOL |  |  |  |  |  |  | 1 | 1 | 4 | 41 | 148 | 57 | 8 | 33 |  | **293** |
| *Eupatorium sp.* | EUPSPP |  |  |  |  |  |  |  | 11 |  | 500 | 3541 | 5708 | 651 | 1220 | 64 | **11695** |
| *Desmodium sp.* | DESSPP |  |  |  |  |  |  |  | 10 | 31 | 106 | 80 | 31 | 16 | 15 | 2 | **291** |
| *Eriochloa villosa* | ERIVIL |  |  |  |  |  |  |  | 6 | 23 | 10 | 2 | 2 |  |  |  | **43** |
| *Sorghastrum nutans* | SORNUT |  |  |  |  |  |  |  | 3 | 57 | 358 | 777 | 974 | 301 | 368 | 14 | **2852** |
| *Eragrostis spectabilis* | ERASPE |  |  |  |  |  |  |  | 3 | 13 | 8 | 87 | 90 | 17 | 844 | 2 | **1064** |
| *Chenopodium album* | CHEALB |  |  |  |  |  |  |  | 2 |  |  |  |  |  |  |  | **2** |
| *Euphorbia corollata* | EUPCOR |  |  |  |  |  |  |  | 1 | 6 | 1 | 3 | 2 | 1 |  |  | **14** |
| *Bidens aristosa* | BIDARI |  |  |  |  |  |  |  | 1 | 5 | 34 | 66 | 35 | 4 | 9 |  | **154** |
| *Strophostyles leiosperma* | STRLEI |  |  |  |  |  |  |  | 1 | 3 | 3 |  |  |  |  |  | **7** |
| *Pycnanthemum pilosum* | PYCPIL |  |  |  |  |  |  |  | 1 |  | 7 | 23 | 49 | 13 | 47 | 5 | **145** |
| *Echinacea pallida* | ECHPAL |  |  |  |  |  |  |  | 1 |  |  |  |  |  |  |  | **1** |
| *Chamaecrista fasciculata* | CHAFAS |  |  |  |  |  |  |  |  | 523 | 609 | 142 | 64 | 34 | 29 | 11 | **1412** |
| *Oenothera filiformis* | OENFIL |  |  |  |  |  |  |  |  | 11 | 29 | 5 | 3 | 1 |  |  | **49** |
| *Lespedeza capitata* | LESCAP |  |  |  |  |  |  |  |  |  | 73 | 88 | 99 | 8 | 42 | 4 | **314** |
| *Conyza canadensis* | CONCAN |  |  |  |  |  |  |  |  |  | 41 | 33 | 18 | 2 |  | 1 | **95** |
| *Solidago altissima* | SOLALT |  |  |  |  |  |  |  |  |  | 38 | 395 | 2636 | 881 | 833 | 35 | **4818** |
| *Rudbeckia subtomentosa* | RUDSUB |  |  |  |  |  |  |  |  |  | 33 | 37 | 227 | 76 | 73 | 8 | **454** |
| *Oenothera biennis* | OENBIE |  |  |  |  |  |  |  |  |  | 22 | 165 | 466 | 72 | 306 | 8 | **1039** |
| *Lespedeza cuneata* | LESCUN |  |  |  |  |  |  |  |  |  | 9 | 84 | 159 | 92 | 535 | 106 | **985** |
| *Cirsium altissimum* | CIRALT |  |  |  |  |  |  |  |  |  | 9 | 3 |  | 2 | 2 |  | **16** |
| *Lespedeza virginica* | LESVIR |  |  |  |  |  |  |  |  |  | 8 | 93 | 284 | 67 | 128 | 10 | **590** |
| *Silphium integrifolium var. integrifolium* | SILINT |  |  |  |  |  |  |  |  |  | 5 | 3 |  | 1 |  |  | **9** |
| *Echinochloa crus-galli* | ECHCRU |  |  |  |  |  |  |  |  |  | 5 | 2 | 1 |  |  |  | **8** |
| *Panicum capillare* | PANCAP |  |  |  |  |  |  |  |  |  | 4 | 2 |  |  | 2 |  | **8** |
| *Erechtites hieracifolius* | EREHIE |  |  |  |  |  |  |  |  |  | 2 |  |  |  |  |  | **2** |
| *Tridens flavus* | TRIFLA |  |  |  |  |  |  |  |  |  | 1 | 7 | 1 | 2 |  |  | **11** |
| *Heliopsis helianthoides* | HELHEL |  |  |  |  |  |  |  |  |  | 1 | 1 | 7 |  |  |  | **9** |
| *Sporobolus heterolepis* | SPOHET |  |  |  |  |  |  |  |  |  | 1 | 1 |  | 1 |  |  | **3** |
| *Leucanthemum vulgare* | LEUVUL |  |  |  |  |  |  |  |  |  | 1 |  |  |  |  |  | **1** |
| *Coreopsis tripteris* | CORTRI |  |  |  |  |  |  |  |  |  |  | 36 | 42 | 46 | 43 | 3 | **170** |
| *Lycopus americanus* | LYCAME |  |  |  |  |  |  |  |  |  |  | 33 |  |  |  |  | **33** |
| *Agalinis sp.* | AGASPP |  |  |  |  |  |  |  |  |  |  | 7 | 31 | 59 | 18 |  | **115** |
| *Boltonia asteroides* | BOLAST |  |  |  |  |  |  |  |  |  |  | 7 | 27 | 27 | 54 | 3 | **118** |
| *Hypericum punctatum* | HYPPUN |  |  |  |  |  |  |  |  |  |  | 4 | 20 | 17 | 86 | 6 | **133** |
| *Symphyotrichum sp.* | SYMSPP |  |  |  |  |  |  |  |  |  |  | 2 | 101 | 383 | 2020 | 138 | **2644** |
| *Amaranthus tuberculatus* | AMATUB |  |  |  |  |  |  |  |  |  |  | 1 | 1 |  | 3 | 3 | **8** |
| *Eleocharis sp.* | ELESPP |  |  |  |  |  |  |  |  |  |  | 1 |  |  |  |  | **1** |
| *Ipomoea hederacea* | IPOHED |  |  |  |  |  |  |  |  |  |  | 1 |  |  |  |  | **1** |
| *Salvia azurea* | SALAZU |  |  |  |  |  |  |  |  |  |  | 1 |  |  |  |  | **1** |
| *Solidago rigida* | SOLRIG |  |  |  |  |  |  |  |  |  |  |  | 11 | 20 | 31 |  | **62** |
| *Euthamia gymnospermoides* | EUTGYM |  |  |  |  |  |  |  |  |  |  |  | 10 | 7 | 20 | 1 | **38** |
| *Liatris pycnostachya var. pycnostachya* | LIAPYC |  |  |  |  |  |  |  |  |  |  |  | 3 |  | 1 |  | **4** |
| *Scirpus georgianus* | SCIGEO |  |  |  |  |  |  |  |  |  |  |  | 1 | 4 | 1 |  | **6** |
| *Gentiana sp.* | GENSPP |  |  |  |  |  |  |  |  |  |  |  | 1 | 1 | 22 | 2 | **26** |
| *Eleocharis sp. 1* | ELESPP1 |  |  |  |  |  |  |  |  |  |  |  | 1 |  |  |  | **1** |
| *Symphyotrichum novae-angliae* | SYMNOV |  |  |  |  |  |  |  |  |  |  |  |  | 14 | 5 |  | **19** |
| *Sporobolus compositus* | SPOCOM |  |  |  |  |  |  |  |  |  |  |  |  | 3 | 3 |  | **6** |
| *Daucus carota* | DAUCAR |  |  |  |  |  |  |  |  |  |  |  |  | 2 |  |  | **2** |
| *Baptisia alba var. macrophylla* | BAPALB |  |  |  |  |  |  |  |  |  |  |  |  | 1 | 5 |  | **6** |
| *Cyperus strigosus* | CYPSTR |  |  |  |  |  |  |  |  |  |  |  |  |  | 9 | 1 | **10** |
| *Solidago nemoralis* | SOLNEM |  |  |  |  |  |  |  |  |  |  |  |  |  | 3 | 1 | **4** |
| *Lysimachia lanceolata* | LYSLAN |  |  |  |  |  |  |  |  |  |  |  |  |  |  | 1 | **1** |
| **Total** | **-** | **5294** | **16006** | **542** | **4189** | **4280** | **1909** | **1459** | **1247** | **2584** | **12435** | **18511** | **25002** | **7794** | **18925** | **986** | **121163** |

**Table S5.** Total number of seeds captured at all the remnant prairie (Tucker Prairie). Divide by 50 to get mean number of seeds per turf grass trap. Multiply by 2 to convert to seeds per m^2^ (total trapping area 0.5 m^2^).

| **Scientific Name** | **SPP6** | **Jun. 13^th^** | **Jun. 27^th^** | **Jul. 10^th^** | **Jul. 26^th^** | **Aug. 8^th^** | **Aug. 22^nd^** | **Sep. 5^th^** | **Sep. 19^th^** | **Oct. 3^rd^** | **Oct. 17^th^** | **Nov. 2^nd^** | **Nov. 14^th^** | **Dec. 1^st^** | **Dec. 12^th^** | **Total** |
| --- | --- | --- | --- | --- | --- | --- | --- | --- | --- | --- | --- | --- | --- | --- | --- | --- |
| *Agrostis hyemalis var. hyemalis* | AGRHYE | 434 | 1557 | 73 | 16 |  | 1 | 3 |  |  |  | 2 |  |  |  | **2086** |
| *Sphenopholis obtusata* | SPHOBT | 147 | 93 | 10 | 4 |  |  | 1 |  |  |  | 1 |  |  |  | **256** |
| *Poa pratensis* | POAPRA | 89 | 180 | 29 | 73 | 15 | 12 | 6 | 2 |  | 2 |  | 2 | 2 |  | **412** |
| *Silene antirrhina* | SILANT | 65 | 9 |  |  |  |  |  |  |  |  |  |  |  |  | **74** |
| *Juncus sp.* | JUNSPP | 4 | 3885 | 340 | 34 | 31 |  | 32 |  |  |  |  |  |  |  | **4326** |
| *Carex festucacea* | CARFES | 4 | 138 | 18 | 7 | 16 | 3 | 4 | 4 | 2 | 2 | 3 | 2 | 1 | 1 | **205** |
| *Plantago virginica* | PLAVIR | 4 | 17 |  |  |  |  |  |  |  |  |  |  |  |  | **21** |
| *Oxalis dillenii* | OXADIL | 2 |  |  |  |  |  |  |  |  |  |  |  |  |  | **2** |
| *Dichanthelium lanuginosum* | DICLAN | 1 | 66 | 25 | 14 | 14 | 11 | 10 | 9 | 3 | 9 | 9 | 5 | 8 | 3 | **187** |
| *Myosotis verna* | MYOVER | 1 | 5 |  |  |  |  |  |  |  |  |  |  |  |  | **6** |
| *Hypoxis hirsuta* | HYPHIR | 1 |  |  |  |  |  |  |  |  |  |  |  |  |  | **1** |
| *Carex bushii* | CARBUS |  | 46 | 23 | 22 | 3 | 10 | 7 | 3 | 1 | 1 | 1 | 1 | 1 |  | **119** |
| *Vulpia octoflora* | VULOCT |  | 20 |  | 3 | 1 |  |  |  |  |  | 1 |  |  |  | **25** |
| *Triodanis perfoliata* | TRIPER |  | 18 | 3 |  |  |  |  |  |  |  |  |  |  |  | **21** |
| *Lobelia spicata* | LOBSPI |  | 16 |  | 1 |  |  |  |  |  |  |  |  |  |  | **17** |
| *Rubus sp.* | RUBSPP |  | 4 | 2 |  | 34 |  |  |  |  | 1 | 1 |  | 1 |  | **43** |
| *Carex bicknellii* | CARBIC |  | 3 |  |  |  |  |  |  |  |  |  |  |  |  | **3** |
| *Pycnanthemum tenuifolium* | PYCTEN |  | 2 | 3 | 1 |  | 2 | 35 | 182 | 797 | 717 | 2187 | 400 | 2330 | 63 | **6719** |
| *Vernonia sp.* | VERSPP |  | 2 |  |  |  |  |  | 1 | 16 | 36 | 26 | 83 | 16 | 15 | **195** |
| *Unknown spp* | UNKSPP |  | 1 | 1 | 1 |  |  | 4 | 1 |  | 1 | 1 | 1 |  | 1 | **12** |
| *Penstemon digitalis* | PENDIG |  | 1 |  |  |  |  |  |  |  |  |  | 7 | 1 |  | **9** |
| *Cardamine sp.* | CARDSP |  | 1 |  |  |  |  |  |  |  |  |  |  |  |  | **1** |
| *Viola sagittata* | VIOSAG |  | 1 |  |  |  |  |  |  |  |  |  |  |  |  | **1** |
| *Erigeron sp.* | ERISPP |  |  | 8 | 3 |  | 2 | 2 | 12 |  |  |  |  |  |  | **27** |
| *Polygala sp.* | POLSPP |  |  | 3 |  | 3 | 2 | 7 | 1 | 1 | 1 |  |  |  |  | **18** |
| *Barbarea vulgaris* | BARVUL |  |  | 2 | 11 | 6 | 5 | 8 | 3 | 12 | 9 | 97 | 8 | 38 | 3 | **202** |
| *Schizachyrium scoparium* | SCHSCO |  |  | 2 | 1 | 2 |  | 1 | 1 | 89 | 188 | 228 | 35 | 80 | 26 | **653** |
| *Setaria parviflora* | SETPAR |  |  |  | 4 | 1 | 8 |  | 4 | 4 | 6 | 6 | 1 | 4 | 2 | **40** |
| *Linum sulcatum var. sulcatum* | LINSUL |  |  |  | 3 | 7 | 3 |  |  |  | 4 | 16 | 5 | 15 | 4 | **57** |
| *Galium obtusum subsp. obtusum* | GALOBT |  |  |  | 3 | 4 | 6 | 9 | 4 |  | 1 |  | 4 | 6 |  | **37** |
| *Achillea millefolium* | ACHMIL |  |  |  | 2 | 19 | 36 | 1 |  | 16 | 3 | 13 | 6 | 8 | 1 | **105** |
| *Koeleria macrantha* | KOEMAC |  |  |  | 2 |  |  |  |  |  |  |  |  |  |  | **2** |
| *Ambrosia artemisiifolia* | AMBART |  |  |  | 1 |  |  |  | 1 | 1 |  |  | 1 |  |  | **4** |
| *Lepidium virginicum* | LEPVIR |  |  |  | 1 |  |  |  |  |  |  |  |  |  |  | **1** |
| *Scirpus pendulus* | SCIPEN |  |  |  | 1 |  |  |  |  |  |  |  |  |  |  | **1** |
| *Crotalaria sagittalis* | CROSAG |  |  |  |  | 2 | 30 | 28 | 48 | 37 | 24 | 11 | 4 | 1 | 2 | **187** |
| *Festuca paradoxa* | FESPAR |  |  |  |  |  | 7 | 3 | 3 | 153 | 5 | 10 | 1 | 2 | 5 | **189** |
| *Helianthus mollis* | HELMOL |  |  |  |  |  | 1 | 1 | 4 | 40 | 58 | 54 | 6 | 33 |  | **197** |
| *Andropogon gerardii* | ANDGER |  |  |  |  |  |  | 23 |  | 53 | 84 | 223 | 24 | 12 | 2 | **421** |
| *Desmodium sp.* | DESSPP |  |  |  |  |  |  | 10 | 20 | 99 | 70 | 29 | 15 | 13 |  | **256** |
| *Rudbeckia hirta var. pulcherrima* | RUDHIR |  |  |  |  |  |  | 3 | 8 | 13 | 15 | 30 | 6 | 9 | 2 | **86** |
| *Acalypha virginica* | ACAVIR |  |  |  |  |  |  | 2 | 5 | 7 | 3 | 3 |  | 1 |  | **21** |
| *Euphorbia corollata* | EUPCOR |  |  |  |  |  |  | 1 | 6 | 1 | 3 | 2 | 1 |  |  | **14** |
| *Chenopodium album* | CHEALB |  |  |  |  |  |  | 1 |  |  |  |  |  |  |  | **1** |
| *Chamaecrista fasciculata* | CHAFAS |  |  |  |  |  |  |  | 41 | 142 | 49 | 9 | 3 |  |  | **244** |
| *Cerastium sp.* | CERSPP |  |  |  |  |  |  |  | 9 |  |  |  |  |  |  | **9** |
| *Sorghastrum nutans* | SORNUT |  |  |  |  |  |  |  | 5 | 72 | 149 | 173 | 30 | 29 | 5 | **463** |
| *Oenothera filiformis* | OENFIL |  |  |  |  |  |  |  | 5 | 4 | 1 |  | 1 |  |  | **11** |
| *Digitaria ischaemum* | DIGISC |  |  |  |  |  |  |  | 2 |  |  |  |  |  |  | **2** |
| *Lespedeza capitata* | LESCAP |  |  |  |  |  |  |  |  | 37 | 47 | 17 | 2 | 3 |  | **106** |
| *Solidago altissima* | SOLALT |  |  |  |  |  |  |  |  | 36 | 282 | 834 | 58 | 199 | 10 | **1419** |
| *Oenothera biennis* | OENBIE |  |  |  |  |  |  |  |  | 20 | 124 | 403 | 50 | 301 | 6 | **904** |
| *Cirsium altissimum* | CIRALT |  |  |  |  |  |  |  |  | 8 | 3 |  | 2 | 2 |  | **15** |
| *Conyza canadensis* | CONCAN |  |  |  |  |  |  |  |  | 5 |  | 3 |  |  | 1 | **9** |
| *Eryngium yuccifolium* | ERYYUC |  |  |  |  |  |  |  |  | 1 | 3 | 28 | 5 | 21 |  | **58** |
| *Sporobolus heterolepis* | SPOHET |  |  |  |  |  |  |  |  | 1 | 1 |  |  |  |  | **2** |
| *Melilotus sp.* | MELSPP |  |  |  |  |  |  |  |  | 1 |  |  | 1 |  |  | **2** |
| *Amorpha canescens* | AMOCAN |  |  |  |  |  |  |  |  | 1 |  |  |  |  |  | **1** |
| *Lycopus americanus* | LYCAME |  |  |  |  |  |  |  |  |  | 33 |  |  |  |  | **33** |
| *Agalinis sp.* | AGASPP |  |  |  |  |  |  |  |  |  | 5 | 27 | 58 | 17 |  | **107** |
| *Tridens flavus* | TRIFLA |  |  |  |  |  |  |  |  |  | 1 |  | 1 |  |  | **2** |
| *Eleocharis sp.* | ELESPP |  |  |  |  |  |  |  |  |  | 1 |  |  |  |  | **1** |
| *Euthamia gymnospermoides* | EUTGYM |  |  |  |  |  |  |  |  |  |  | 10 | 7 | 20 | 1 | **38** |
| *Symphyotrichum sp.* | SYMSPP |  |  |  |  |  |  |  |  |  |  | 4 | 8 | 45 | 3 | **60** |
| *Bidens aristosa* | BIDARI |  |  |  |  |  |  |  |  |  |  | 3 | 3 |  |  | **6** |
| *Eupatorium sp.* | EUPSPP |  |  |  |  |  |  |  |  |  |  | 2 | 28 |  |  | **30** |
| *Monarda fistulosa subsp. Fistulosa* | MONFIS |  |  |  |  |  |  |  |  |  |  | 2 | 1 |  |  | **3** |
| *Ratibida pinnata* | RATPIN |  |  |  |  |  |  |  |  |  |  | 1 | 5 |  |  | **6** |
| *Hypericum punctatum* | HYPPUN |  |  |  |  |  |  |  |  |  |  | 1 |  |  |  | **1** |
| *Rudbeckia subtomentosa* | RUDSUB |  |  |  |  |  |  |  |  |  |  |  | 4 |  |  | **4** |
| *Scirpus georgianus* | SCIGEO |  |  |  |  |  |  |  |  |  |  |  | 4 |  |  | **4** |
| *Coreopsis tripteris* | CORTRI |  |  |  |  |  |  |  |  |  |  |  | 2 |  |  | **2** |
| *Eragrostis spectabilis* | ERASPE |  |  |  |  |  |  |  |  |  |  |  |  | 1 |  | **1** |
| *Gentiana sp.* | GENSPP |  |  |  |  |  |  |  |  |  |  |  |  | 1 |  | **1** |
| *Lespedeza cuneata* | LESCUN |  |  |  |  |  |  |  |  |  |  |  |  | 1 |  | **1** |
| *Verbena hastata* | VERHAS |  |  |  |  |  |  |  |  |  |  |  |  | 1 |  | **1** |
| *Lysimachia lanceolata* | LYSLAN |  |  |  |  |  |  |  |  |  |  |  |  |  | 1 | **1** |
| Total | - | **752** | **6065** | **542** | **208** | **158** | **139** | **202** | **384** | **1673** | **1942** | **4471** | **891** | **3223** | **157** | **20807** |

**Table S6.** Total number of seeds captured at all the young restored prairie (2-year-old). Divide by 50 to get mean number of seeds per turf grass trap. Multiply by 2 to convert to seeds per m^2^ (total trapping area 0.5 m^2^).

| **Scientific Name** | **SPP6** | **Ju. 13^th^** | **Jun. 27^th^** | **Jul. 11^th^** | **Jul. 26^th^** | **Aug. 8^th^** | **Aug. 22^nd^** | **Sep. 5^th^** | **Sep. 19^th^** | **Oct. 3^rd^** | **Oct. 17^th^** | **Nov. 2^nd^** | **Nov. 14^th^** | **Dec. 1^st^** | **Dec. 12^th^** | **Total** |
| --- | --- | --- | --- | --- | --- | --- | --- | --- | --- | --- | --- | --- | --- | --- | --- | --- |
| *Juncus sp.* | JUNSPP | 233 | 318 | 52 | 34 |  |  | 2 |  |  |  |  |  |  |  | **639** |
| *Cerastium sp.* | CERSPP | 149 | 61 | 11 | 3 | 6 |  |  |  |  |  |  |  |  | 4 | **234** |
| *Veronica sp.* | VEROSP | 102 | 69 | 14 | 5 | 3 | 3 |  |  | 2 |  |  |  |  |  | **198** |
| *Myosotis verna* | MYOVER | 64 | 44 | 11 | 4 | 4 | 1 | 5 |  | 1 |  | 3 |  | 1 | 1 | **139** |
| *Plantago virginica* | PLAVIR | 50 | 58 | 38 | 5 | 1 | 1 |  |  |  |  | 1 | 1 | 1 | 1 | **157** |
| *Triodanis perfoliata* | TRIPER | 24 | 4 | 7 | 13 |  |  |  |  |  |  |  |  |  |  | **48** |
| *Sphenopholis obtusata* | SPHOBT | 14 | 6 | 1 |  |  |  | 1 |  | 1 | 1 |  |  |  |  | **24** |
| *Geranium carolinianum* | GERCAR | 11 | 15 | 2 |  |  |  |  |  |  |  |  |  |  |  | **28** |
| *Capsella bursa-pastoris* | CAPBUR | 5 | 12 |  | 1 |  |  |  |  |  |  |  |  |  |  | **18** |
| *Alopecurus carolinianus* | ALOCAR | 3 |  |  |  |  |  |  |  |  |  |  |  |  |  | **3** |
| *Lepidium virginicum* | LEPVIR | 2 | 10 | 1 |  |  |  |  |  |  |  |  |  |  |  | **13** |
| *Galium aparine* | GALAPA | 2 | 9 |  | 1 |  |  |  |  |  |  |  |  |  |  | **12** |
| *Thlaspi arvense* | THLARV | 2 | 4 |  |  |  | 1 | 3 |  |  |  | 4 | 1 | 1 |  | **16** |
| *Oxalis dillenii* | OXADIL | 2 | 1 | 1 | 3 | 1 | 2 |  |  |  |  | 3 |  | 1 |  | **14** |
| *Poa pratensis* | POAPRA | 1 | 7 | 1 | 1 | 10 | 6 |  |  |  |  |  |  |  |  | **26** |
| *Dichanthelium lanuginosum* | DICLAN | 1 | 2 | 1 |  |  |  |  | 10 |  |  |  |  |  |  | **14** |
| *Melilotus sp.* | MELSPP | 1 |  |  |  |  | 2 | 1 | 9 | 1 |  |  | 1 |  |  | **15** |
| *Platanus occidentalis* | PLAOCC | 1 |  |  |  |  |  |  |  |  |  |  |  |  |  | **1** |
| *Erigeron sp.* | ERISPP |  | 78 | 1876 | 2483 | 1290 | 625 | 58 | 63 | 40 |  |  |  |  |  | **6513** |
| *Digitaria ischaemum* | DIGISC |  | 22 | 2 | 1 | 6 | 3 | 1 | 257 | 3637 | 7585 | 3643 | 883 | 556 | 163 | **16759** |
| *Setaria sp.* | SETSPP |  | 16 |  |  | 1 | 5 | 16 | 208 | 1794 | 1484 | 979 | 107 | 188 | 24 | **4822** |
| *Anagallis minima* | ANAMIN |  | 15 | 35 | 4 | 1 |  |  |  |  |  |  |  |  |  | **55** |
| *Barbarea vulgaris* | BARVUL |  | 7 |  |  |  | 1 |  |  |  |  |  |  |  |  | **8** |
| *Ambrosia artemisiifolia* | AMBART |  | 2 |  |  | 1 |  | 4 |  | 15 | 25 | 13 | 3 | 8 | 2 | **73** |
| *Medicago lupulina* | MEDLUP |  | 1 |  | 1 |  | 2 | 11 | 2 |  | 1 |  |  |  |  | **18** |
| *Kummerowia sp.* | KUMSPP |  | 1 |  |  |  | 1 | 4 | 1 | 20 | 41 | 70 | 56 | 118 | 12 | **324** |
| *Verbena hastata* | VERHAS |  | 1 |  |  |  |  |  |  |  |  | 4 |  | 4 | 1 | **10** |
| *Koeleria macrantha* | KOEMAC |  | 1 |  |  |  |  |  |  |  |  |  |  |  |  | **1** |
| *Taraxacum officinale* | TAROFF |  | 1 |  |  |  |  |  |  |  |  |  |  |  |  | **1** |
| *Scleria triglomerata* | SCLTRI |  |  | 4 |  |  |  |  |  |  |  |  |  |  |  | **4** |
| *Coreopsis lanceolata* | CORLAN |  |  | 2 | 3 |  | 1 |  |  | 1 |  |  |  |  |  | **7** |
| *Festuca arundinacea* | FESARU |  |  | 1 |  |  |  | 1 |  |  |  |  |  |  |  | **2** |
| *Carex bushii* | CARBUS |  |  | 1 |  |  |  |  |  |  |  |  | 1 |  |  | **2** |
| *Achillea millefolium* | ACHMIL |  |  |  | 13 | 3 | 5 | 23 | 19 | 15 | 25 | 49 | 41 | 26 | 5 | **224** |
| *Unknown spp* | UNKSPP |  |  |  | 1 |  | 1 | 1 | 2 |  |  | 2 |  |  |  | **7** |
| *Bromus japonicus* | BROJAP |  |  |  |  | 27 | 5 |  | 4 |  |  |  |  | 21 | 1 | **58** |
| *Cyperus echinatus* | CYPECH |  |  |  |  | 21 | 174 | 108 | 11 | 12 | 24 | 24 | 22 | 19 | 7 | **422** |
| *Acalypha virginica* | ACAVIR |  |  |  |  | 2 | 1 | 5 | 21 | 38 | 16 | 12 |  | 1 |  | **96** |
| *Rudbeckia hirta var. pulcherrima* | RUDHIR |  |  |  |  | 1 |  | 1 | 4 | 20 | 41 | 75 | 28 | 54 | 5 | **229** |
| *Ratibida pinnata* | RATPIN |  |  |  |  |  | 1 | 15 | 11 | 137 | 78 | 117 | 48 | 89 | 5 | **501** |
| *Pycnanthemum tenuifolium* | PYCTEN |  |  |  |  |  | 1 |  |  |  | 1 | 1 | 1 |  |  | **4** |
| *Eupatorium sp.* | EUPSPP |  |  |  |  |  |  | 11 |  | 446 | 3453 | 5553 | 604 | 1196 | 63 | **11326** |
| *Eriochloa villosa* | ERIVIL |  |  |  |  |  |  | 6 | 23 | 10 | 2 | 2 |  |  |  | **43** |
| *Penstemon digitalis* | PENDIG |  |  |  |  |  |  | 2 | 42 | 894 | 775 | 3462 | 1064 | 5298 | 100 | **11637** |
| *Festuca paradoxa* | FESPAR |  |  |  |  |  |  | 1 | 4 | 3 | 5 | 1 | 2 |  |  | **16** |
| *Strophostyles leiosperma* | STRLEI |  |  |  |  |  |  | 1 | 1 |  |  |  |  |  |  | **2** |
| *Chenopodium album* | CHEALB |  |  |  |  |  |  | 1 |  |  |  |  |  |  |  | **1** |
| *Chamaecrista fasciculata* | CHAFAS |  |  |  |  |  |  |  | 468 | 431 | 87 | 52 | 31 | 29 | 11 | **1109** |
| *Desmodium sp.* | DESSPP |  |  |  |  |  |  |  | 4 |  | 4 | 1 | 1 |  | 1 | **11** |
| *Bidens aristosa* | BIDARI |  |  |  |  |  |  |  | 3 | 30 | 61 | 24 | 1 | 4 |  | **123** |
| *Vernonia sp.* | VERSPP |  |  |  |  |  |  |  | 2 |  | 2 |  |  |  |  | **4** |
| *Sorghastrum nutans* | SORNUT |  |  |  |  |  |  |  | 1 | 19 | 24 | 25 | 15 | 8 |  | **92** |
| *Conyza canadensis* | CONCAN |  |  |  |  |  |  |  |  | 32 | 32 | 13 | 1 |  |  | **78** |
| *Schizachyrium scoparium* | SCHSCO |  |  |  |  |  |  |  |  | 5 | 1 | 1 | 1 | 1 |  | **9** |
| *Panicum capillare* | PANCAP |  |  |  |  |  |  |  |  | 4 | 2 |  |  |  |  | **6** |
| *Echinochloa crus-galli* | ECHCRU |  |  |  |  |  |  |  |  | 3 | 1 | 1 |  |  |  | **5** |
| *Solidago altissima* | SOLALT |  |  |  |  |  |  |  |  | 2 | 85 | 1426 | 745 | 577 | 23 | **2858** |
| *Rudbeckia subtomentosa* | RUDSUB |  |  |  |  |  |  |  |  | 2 | 13 | 16 | 6 | 2 |  | **39** |
| *Erechtites hieracifolius* | EREHIE |  |  |  |  |  |  |  |  | 2 |  |  |  |  |  | **2** |
| *Silphium integrifolium var. integrifolium* | SILINT |  |  |  |  |  |  |  |  | 1 | 3 |  | 1 |  |  | **5** |
| *Oenothera filiformis* | OENFIL |  |  |  |  |  |  |  |  | 1 | 1 |  |  |  |  | **2** |
| *Lespedeza capitata* | LESCAP |  |  |  |  |  |  |  |  | 1 |  | 5 |  | 4 |  | **10** |
| *Cirsium altissimum* | CIRALT |  |  |  |  |  |  |  |  | 1 |  |  |  |  |  | **1** |
| *Leucanthemum vulgare* | LEUVUL |  |  |  |  |  |  |  |  | 1 |  |  |  |  |  | **1** |
| *Vulpia octoflora* | VULOCT |  |  |  |  |  |  |  |  | 1 |  |  |  |  |  | **1** |
| *Eragrostis spectabilis* | ERASPE |  |  |  |  |  |  |  |  |  | 84 | 52 | 3 | 52 | 1 | **192** |
| *Oenothera biennis* | OENBIE |  |  |  |  |  |  |  |  |  | 41 | 62 | 22 | 3 | 1 | **129** |
| *Coreopsis tripteris* | CORTRI |  |  |  |  |  |  |  |  |  | 26 | 38 | 36 | 41 | 3 | **144** |
| *Lespedeza cuneata* | LESCUN |  |  |  |  |  |  |  |  |  | 20 | 64 | 13 | 166 | 6 | **269** |
| *Hypericum punctatum* | HYPPUN |  |  |  |  |  |  |  |  |  | 4 | 19 | 17 | 85 | 6 | **131** |
| *Agalinis sp.* | AGASPP |  |  |  |  |  |  |  |  |  | 2 | 1 |  |  |  | **3** |
| *Tridens flavus* | TRIFLA |  |  |  |  |  |  |  |  |  | 1 | 1 | 1 |  |  | **3** |
| *Monarda fistulosa subsp. Fistulosa* | MONFIS |  |  |  |  |  |  |  |  |  | 1 |  | 1 |  |  | **2** |
| *Symphyotrichum sp.* | SYMSPP |  |  |  |  |  |  |  |  |  |  | 84 | 318 | 1827 | 127 | **2356** |
| *Carex festucacea* | CARFES |  |  |  |  |  |  |  |  |  |  | 8 |  |  |  | **8** |
| *Pycnanthemum pilosum* | PYCPIL |  |  |  |  |  |  |  |  |  |  | 3 |  |  |  | **3** |
| *Boltonia asteroides* | BOLAST |  |  |  |  |  |  |  |  |  |  | 1 | 2 | 8 |  | **11** |
| *Andropogon gerardii* | ANDGER |  |  |  |  |  |  |  |  |  |  | 1 | 2 |  |  | **3** |
| *Scirpus georgianus* | SCIGEO |  |  |  |  |  |  |  |  |  |  | 1 |  | 1 |  | **2** |
| *Amaranthus tuberculatus* | AMATUB |  |  |  |  |  |  |  |  |  |  | 1 |  |  |  | **1** |
| *Daucus carota* | DAUCAR |  |  |  |  |  |  |  |  |  |  |  | 2 |  |  | **2** |
| *Solidago rigida* | SOLRIG |  |  |  |  |  |  |  |  |  |  |  | 1 | 6 |  | **7** |
| *Symphyotrichum novae-angliae* | SYMNOV |  |  |  |  |  |  |  |  |  |  |  | 1 | 2 |  | **3** |
| *Eryngium yuccifolium* | ERYYUC |  |  |  |  |  |  |  |  |  |  |  | 1 |  |  | **1** |
| *Cyperus strigosus* | CYPSTR |  |  |  |  |  |  |  |  |  |  |  |  | 9 | 1 | **10** |
| *Liatris pycnostachya var. pycnostachya* | LIAPYC |  |  |  |  |  |  |  |  |  |  |  |  | 1 |  | **1** |
| *Mollugo verticillata* | MOLVER |  |  |  |  |  |  |  |  |  |  |  |  | 1 |  | **1** |
| *Lespedeza virginica* | LESVIR |  |  |  |  |  |  |  |  |  |  |  |  |  | 1 | **1** |
| **Total** | **-** | **667** | **765** | **2061** | **2576** | **1378** | **842** | **282** | **1170** | **7623** | **14052** | **15918** | **4085** | **10409** | **575** | **62403** |

**Table S6.** Total number of seeds captured at all the middle-aged prairie (5-6-year-old). Divide by 50 to get mean number of seeds per turf grass trap. Multiply by 2 to convert to seeds per m^2^ (total trapping area 0.5 m^2^).

| **Scientific Name** | **SPP6** | **Jun. 13^th^** | **Jun. 27^th^** | **Jul. 11^th^** | **Jul. 26^th^** | **Aug. 08^th^** | **Aug. 22^nd^** | **Sep. 5^th^** | **Sep. 19^th^** | **Oct. 3th** | **Oct. 17^th^** | **Nov. 2^nd^** | **Nov. 14^th^** | **Dec. 1^st^** | **Dec. 12^th^** | **Total** |
| --- | --- | --- | --- | --- | --- | --- | --- | --- | --- | --- | --- | --- | --- | --- | --- | --- |
| *Sphenopholis obtusata* | SPHOBT | 1159 | 863 | 72 | 116 | 5 | 2 | 38 | 8 |  | 3 | 11 |  | 45 | 1 | **2323** |
| *Coreopsis lanceolata* | CORLAN | 143 | 53 | 46 | 39 | 1 | 7 | 28 | 15 | 4 | 7 | 9 |  | 6 |  | **358** |
| *Plantago virginica* | PLAVIR | 42 | 22 | 6 | 9 | 1 | 3 | 5 |  | 7 | 5 | 8 | 7 | 9 | 1 | **125** |
| *Juncus sp.* | JUNSPP | 27 | 2511 | 512 | 189 | 51 | 10 | 73 |  |  |  |  |  | 98 |  | **3471** |
| *Cerastium sp.* | CERSPP | 9 | 10 | 1 | 1 |  | 1 | 1 | 1 |  |  |  |  |  |  | **24** |
| *Poa pratensis* | POAPRA | 5 | 51 | 298 | 44 | 12 | 8 | 25 | 28 | 2 | 4 | 5 |  | 1 | 1 | **484** |
| *Veronica sp.* | VEROSP | 5 | 8 | 2 | 2 | 2 | 3 | 2 | 3 | 1 |  |  |  |  |  | **28** |
| *Myosotis verna* | MYOVER | 4 | 4 |  | 1 |  |  | 1 |  |  |  |  |  |  |  | **10** |
| *Hordeum pusillum* | HORPUS | 3 | 7 |  |  |  |  |  |  |  |  |  |  |  |  | **10** |
| *Ratibida pinnata* | RATPIN | 2 | 1 | 1 | 12 |  | 1 | 21 | 27 | 20 | 41 | 45 | 9 | 47 |  | **227** |
| *Geranium carolinianum* | GERCAR | 2 |  |  |  |  |  |  |  |  |  |  |  |  |  | **2** |
| *Carex bushii* | CARBUS | 1 | 12 | 7 | 9 | 1 | 1 | 4 | 2 |  | 1 | 2 | 1 | 2 | 1 | **44** |
| *Unknown spp* | UNKSPP | 1 | 2 |  | 1 |  | 1 | 4 |  | 2 |  |  |  |  |  | **11** |
| *Galium aparine* | GALAPA | 1 |  |  |  |  |  |  |  |  |  |  |  |  |  | **1** |
| *Erigeron sp.* | ERISPP |  | 70 | 528 | 432 | 84 | 66 | 3 | 7 |  |  |  |  |  |  | **1190** |
| *Monarda fistulosa subsp. Fistulosa* | MONFIS |  | 18 |  | 4 | 4 | 8 | 35 | 32 | 107 | 24 | 44 | 16 | 64 |  | **356** |
| *Setaria sp.* | SETSPP |  | 14 | 11 |  |  |  | 7 | 60 | 275 | 205 | 115 | 70 | 54 |  | **811** |
| *Coreopsis palmata* | CORPAL |  | 13 | 17 | 3 | 1 |  |  |  |  |  |  |  |  |  | **34** |
| *Festuca arundinacea* | FESARU |  | 9 | 2 |  | 1 |  |  |  |  |  |  |  |  |  | **12** |
| *Carex cephalophora* | CARCEP |  | 8 |  |  |  |  |  |  |  |  |  |  |  |  | **8** |
| *Medicago lupulina* | MEDLUP |  | 6 | 52 | 95 | 65 | 76 | 50 | 24 | 7 | 7 |  | 1 | 5 |  | **388** |
| *Bromus japonicus* | BROJAP |  | 4 | 5 | 2 | 1 |  | 5 |  |  |  |  |  | 4 |  | **21** |
| *Barbarea vulgaris* | BARVUL |  | 4 | 2 | 1 |  |  | 3 | 2 |  | 2 |  | 3 | 7 | 1 | **25** |
| *Oxalis dillenii* | OXADIL |  | 2 |  |  |  | 1 |  |  |  | 1 | 5 | 3 |  |  | **12** |
| *Rumex crispus* | RUMCRI |  | 2 |  |  |  |  |  |  |  |  |  |  |  |  | **2** |
| *Pycnanthemum tenuifolium* | PYCTEN |  | 1 |  | 5 | 2 | 2 | 8 | 215 | 230 | 92 | 453 | 306 | 356 | 37 | **1707** |
| *Eryngium yuccifolium* | ERYYUC |  | 1 |  | 1 |  |  |  |  | 4 | 14 | 32 | 21 | 14 | 3 | **90** |
| *Persicaria longiseta* | PERLON |  | 1 |  |  |  |  |  |  |  |  |  |  |  |  | **1** |
| *Koeleria macrantha* | KOEMAC |  |  | 65 | 107 | 17 | 10 | 58 | 7 |  |  |  |  | 1 |  | **265** |
| *Melilotus sp.* | MELSPP |  |  | 22 | 16 | 22 | 103 | 79 | 119 | 50 | 15 | 14 | 8 | 6 |  | **454** |
| *Anagallis minima* | ANAMIN |  |  | 17 | 9 |  |  | 2 | 9 |  |  |  |  |  |  | **37** |
| *Scirpus pendulus* | SCIPEN |  |  | 7 | 16 |  |  |  |  |  | 1 | 5 | 1 | 2 |  | **32** |
| *Blephilia ciliata* | BLECIL |  |  | 2 |  |  |  |  |  |  |  |  |  |  |  | **2** |
| *Lepidium virginicum* | LEPVIR |  |  | 1 |  |  |  |  |  |  |  |  |  |  |  | **1** |
| *Rudbeckia hirta var. pulcherrima* | RUDHIR |  |  |  | 36 |  |  | 9 | 11 | 99 | 22 | 45 | 41 | 34 | 14 | **311** |
| *Cyperus acuminatus* | CYPACU |  |  |  | 18 |  | 8 | 25 | 1 |  |  | 3 |  | 3 |  | **58** |
| *Festuca paradoxa* | FESPAR |  |  |  | 4 | 5 | 2 | 7 | 11 | 5 | 3 | 1 | 1 | 5 |  | **44** |
| *Cyperus echinatus* | CYPECH |  |  |  | 3 | 1 | 7 | 6 | 6 | 1 | 7 | 6 | 1 | 11 |  | **49** |
| *Lythrum alatum* | LYTALA |  |  |  | 2 |  |  |  |  |  |  |  |  |  |  | **2** |
| *Ambrosia artemisiifolia* | AMBART |  |  |  | 1 |  |  |  |  | 2 | 1 |  |  |  |  | **4** |
| *Dianthus armeria* | DIAARM |  |  |  |  | 1 |  |  |  |  |  |  |  | 2 |  | **3** |
| *Penstemon digitalis* | PENDIG |  |  |  |  |  | 6 | 6 | 36 | 322 | 158 | 772 | 1249 | 1757 | 21 | **4327** |
| *Digitaria ischaemum* | DIGISC |  |  |  |  |  | 2 | 13 | 210 | 1097 | 421 | 195 | 72 | 36 | 2 | **2048** |
| *Eragrostis spectabilis* | ERASPE |  |  |  |  |  |  | 3 | 13 | 8 | 3 | 38 | 14 | 778 | 1 | **858** |
| *Sorghastrum nutans* | SORNUT |  |  |  |  |  |  | 2 | 39 | 127 | 362 | 539 | 149 | 223 | 5 | **1446** |
| *Acalypha virginica* | ACAVIR |  |  |  |  |  |  | 2 | 1 | 4 | 5 | 1 |  |  |  | **13** |
| *Schizachyrium scoparium* | SCHSCO |  |  |  |  |  |  | 1 | 14 | 79 | 24 | 35 | 18 | 15 | 2 | **188** |
| *Bidens aristosa* | BIDARI |  |  |  |  |  |  | 1 | 2 | 4 | 5 | 8 |  | 5 |  | **25** |
| *Pycnanthemum pilosum* | PYCPIL |  |  |  |  |  |  | 1 |  | 7 | 23 | 46 | 13 | 44 | 2 | **136** |
| *Thlaspi arvense* | THLARV |  |  |  |  |  |  | 1 |  |  |  |  | 1 | 1 | 1 | **4** |
| *Dichanthelium lanuginosum* | DICLAN |  |  |  |  |  |  | 1 |  |  |  |  |  |  |  | **1** |
| *Echinacea pallida* | ECHPAL |  |  |  |  |  |  | 1 |  |  |  |  |  |  |  | **1** |
| *Desmodium sp.* | DESSPP |  |  |  |  |  |  |  | 2 | 1 |  | 1 |  | 1 |  | **5** |
| *Andropogon gerardii* | ANDGER |  |  |  |  |  |  |  | 2 |  | 28 | 52 | 4 | 24 | 1 | **111** |
| *Chamaecrista fasciculata* | CHAFAS |  |  |  |  |  |  |  | 1 | 5 | 1 |  |  |  |  | **7** |
| *Kummerowia sp.* | KUMSPP |  |  |  |  |  |  |  | 1 | 2 | 5 | 4 |  | 12 |  | **24** |
| *Eupatorium sp.* | EUPSPP |  |  |  |  |  |  |  |  | 54 | 88 | 153 | 18 | 23 | 1 | **337** |
| *Rudbeckia subtomentosa* | RUDSUB |  |  |  |  |  |  |  |  | 17 | 12 | 175 | 44 | 50 | 6 | **304** |
| *Lespedeza capitata* | LESCAP |  |  |  |  |  |  |  |  | 14 | 4 | 1 | 1 | 2 | 1 | **23** |
| *Lespedeza cuneata* | LESCUN |  |  |  |  |  |  |  |  | 9 | 34 | 66 | 44 | 101 | 5 | **259** |
| *Lespedeza virginica* | LESVIR |  |  |  |  |  |  |  |  | 4 | 5 | 5 | 6 | 1 | 1 | **22** |
| *Conyza canadensis* | CONCAN |  |  |  |  |  |  |  |  | 3 | 1 | 2 | 1 |  |  | **7** |
| *Oenothera biennis* | OENBIE |  |  |  |  |  |  |  |  | 2 |  | 1 |  | 2 | 1 | **6** |
| *Silphium integrifolium var. integrifolium* | SILINT |  |  |  |  |  |  |  |  | 2 |  |  |  |  |  | **2** |
| *Helianthus mollis* | HELMOL |  |  |  |  |  |  |  |  | 1 | 90 | 2 | 2 |  |  | **95** |
| *Tridens flavus* | TRIFLA |  |  |  |  |  |  |  |  | 1 | 2 |  |  |  |  | **3** |
| *Echinochloa crus-galli* | ECHCRU |  |  |  |  |  |  |  |  | 1 | 1 |  |  |  |  | **2** |
| *Solidago altissima* | SOLALT |  |  |  |  |  |  |  |  |  | 27 | 286 | 67 | 47 | 1 | **428** |
| *Coreopsis tripteris* | CORTRI |  |  |  |  |  |  |  |  |  | 10 | 4 | 6 | 2 |  | **22** |
| *Boltonia asteroides* | BOLAST |  |  |  |  |  |  |  |  |  | 7 | 26 | 25 | 44 | 3 | **105** |
| *Symphyotrichum sp.* | SYMSPP |  |  |  |  |  |  |  |  |  | 2 | 12 | 38 | 78 | 1 | **131** |
| *Amaranthus tuberculatus* | AMATUB |  |  |  |  |  |  |  |  |  | 1 |  |  | 3 | 3 | **7** |
| *Solidago rigida* | SOLRIG |  |  |  |  |  |  |  |  |  |  | 11 | 19 | 25 |  | **55** |
| *Agalinis sp.* | AGASPP |  |  |  |  |  |  |  |  |  |  | 3 | 1 | 1 |  | **5** |
| *Liatris pycnostachya var. pycnostachya* | LIAPYC |  |  |  |  |  |  |  |  |  |  | 3 |  |  |  | **3** |
| *Eleocharis sp. 1* | ELESPP1 |  |  |  |  |  |  |  |  |  |  | 1 |  |  |  | **1** |
| *Symphyotrichum novae-angliae* | SYMNOV |  |  |  |  |  |  |  |  |  |  |  | 13 | 3 |  | **16** |
| *Sporobolus compositus* | SPOCOM |  |  |  |  |  |  |  |  |  |  |  | 1 | 3 |  | **4** |
| *Linum sulcatum var. sulcatum* | LINSUL |  |  |  |  |  |  |  |  |  |  |  | 1 | 1 |  | **2** |
| *Sporobolus heterolepis* | SPOHET |  |  |  |  |  |  |  |  |  |  |  | 1 |  |  | **1** |
| *Solidago nemoralis* | SOLNEM |  |  |  |  |  |  |  |  |  |  |  |  | 3 |  | **3** |
| *Panicum capillare* | PANCAP |  |  |  |  |  |  |  |  |  |  |  |  | 2 |  | **2** |
| *Hypericum punctatum* | HYPPUN |  |  |  |  |  |  |  |  |  |  |  |  | 1 |  | **1** |
| **Total** | **-** | **1404** | **3697** | **1676** | **1178** | **277** | **328** | **531** | **909** | **2580** | **1774** | **3245** | **2297** | **4064** | **117** | **24077** |

**Table S7.** Total number of seeds captured at all the old prairie (15-year-old). Divide by 50 to get mean number of seeds per turf grass trap. Multiply by 2 to convert to seeds per m^2^ (total trapping area 0.5 m^2^).

| **Scientific Name** | **SPP6** | **Jun. 13^th^** | **Jun. 27^th^** | **Jul. 11^th^** | **Jul. 26^th^** | **Aug. 8^th^** | **Aug. 22^nd^** | **Sep. 5^th^** | **Sep. 19^th^** | **Oct. 3^rd^** | **Oct. 17^th^** | **Nov. 2^nd^** | **Nov. 14^th^** | **Dec. 1^st^** | **Dec. 12^th^** | **Total** |
| --- | --- | --- | --- | --- | --- | --- | --- | --- | --- | --- | --- | --- | --- | --- | --- | --- |
| *Agrostis hyemalis var. hyemalis* | AGRHYE | 1576 | 1103 | 53 | 11 |  | 4 | 2 |  |  | 4 |  |  |  |  | **2753** |
| *Juncus sp.* | JUNSPP | 669 | 4104 | 217 | 155 | 27 | 56 | 28 |  |  |  |  |  |  |  | **5256** |
| *Sphenopholis obtusata* | SPHOBT | 150 | 70 |  | 4 | 2 |  |  |  |  |  |  |  |  |  | **226** |
| *Carex bushii* | CARBUS | 16 | 9 | 20 | 10 | 6 | 2 | 1 | 1 |  | 2 | 4 |  | 2 |  | **73** |
| *Alopecurus carolinianus* | ALOCAR | 14 | 1 | 2 |  |  |  |  |  |  |  |  |  |  |  | **17** |
| *Anagallis minima* | ANAMIN | 11 |  | 15 |  |  |  |  |  |  |  |  |  |  |  | **26** |
| *Carex festucacea* | CARFES | 10 | 84 | 85 | 51 | 11 | 16 | 130 | 12 | 12 | 5 | 17 | 2 | 7 |  | **442** |
| *Poa pratensis* | POAPRA | 10 | 4 | 3 |  |  |  |  |  |  |  |  |  |  |  | **17** |
| *Penstemon digitalis* | PENDIG | 4 |  |  | 35 |  |  |  | 1 | 22 | 58 | 152 | 106 | 277 | 7 | **662** |
| *Tradescantia ohiensis* | TRAOHI | 3 | 66 | 33 | 12 |  | 7 |  |  | 1 |  | 1 | 1 |  |  | **124** |
| *Plantago virginica* | PLAVIR | 2 | 2 | 2 |  |  |  |  |  |  |  |  |  |  |  | **6** |
| *Capsella bursa-pastoris* | CAPBUR | 2 |  |  | 2 |  |  |  |  |  |  |  |  |  |  | **4** |
| *Oxalis dillenii* | OXADIL | 1 | 1 |  |  | 1 |  |  |  | 1 | 2 | 3 | 1 |  |  | **10** |
| *Unknown spp* | UNKSPP | 1 | 1 |  |  |  |  | 1 |  | 1 |  |  |  |  |  | **4** |
| *Eryngium yuccifolium* | ERYYUC | 1 |  | 3 |  | 7 | 1 | 9 | 6 | 46 | 92 | 110 | 23 | 47 | 1 | **346** |
| *Veronica sp.* | VEROSP | 1 |  |  |  |  |  |  |  |  |  |  |  |  |  | **1** |
| *Vulpia octoflora* | VULOCT |  | 14 | 7 | 1 | 1 | 2 |  |  |  |  |  |  |  |  | **25** |
| *Achillea millefolium* | ACHMIL |  | 14 |  |  |  |  |  |  |  |  | 2 |  |  |  | **16** |
| *Myosotis verna* | MYOVER |  | 5 |  |  |  |  |  | 1 |  |  |  |  |  |  | **6** |
| *Lepidium virginicum* | LEPVIR |  | 1 | 1 |  |  |  |  |  |  |  |  |  |  |  | **2** |
| *Erigeron sp.* | ERISPP |  |  | 10 | 6 | 9 | 1 | 1 | 5 |  |  |  |  |  |  | **32** |
| *Andropogon gerardii* | ANDGER |  |  | 1 |  |  |  |  | 1 | 10 | 9 | 11 | 1 | 9 |  | **42** |
| *Carex annectens* | CARANN |  |  |  | 15 |  |  |  |  |  |  |  |  |  |  | **15** |
| *Melilotus sp.* | MELSPP |  |  |  | 6 | 29 | 57 | 45 | 28 | 10 | 7 | 2 |  | 1 |  | **185** |
| *Amorpha canescens* | AMOCAN |  |  |  | 4 | 1 | 1 |  | 1 |  |  | 1 | 2 | 13 |  | **23** |
| *Cyperus acuminatus* | CYPACU |  |  |  | 2 |  |  | 5 | 4 | 12 | 5 | 6 | 4 | 7 |  | **45** |
| *Pycnanthemum tenuifolium* | PYCTEN |  |  |  | 1 | 1 |  | 7 | 10 | 60 | 51 | 85 | 54 | 112 |  | **381** |
| *Medicago lupulina* | MEDLUP |  |  |  | 1 |  |  |  | 9 | 3 |  |  |  |  |  | **13** |
| *Ratibida pinnata* | RATPIN |  |  |  | 1 |  |  |  |  |  | 1 | 4 |  |  |  | **6** |
| *Festuca paradoxa* | FESPAR |  |  |  | 1 |  |  |  |  |  |  |  |  |  |  | **1** |
| *Acalypha virginica* | ACAVIR |  |  |  |  | 1 |  | 1 | 1 |  | 1 |  |  |  |  | **4** |
| *Mollugo verticillata* | MOLVER |  |  |  |  |  | 2 | 1 |  |  |  | 1 |  | 7 |  | **11** |
| *Kummerowia sp.* | KUMSPP |  |  |  |  |  | 1 |  |  |  | 9 | 7 | 3 |  |  | **20** |
| *Sorghastrum nutans* | SORNUT |  |  |  |  |  |  | 1 | 12 | 140 | 242 | 237 | 107 | 108 | 4 | **851** |
| *Chamaecrista fasciculata* | CHAFAS |  |  |  |  |  |  |  | 13 | 31 | 5 | 3 |  |  |  | **52** |
| *Oenothera filiformis* | OENFIL |  |  |  |  |  |  |  | 6 | 24 | 3 | 3 |  |  |  | **36** |
| *Desmodium sp.* | DESSPP |  |  |  |  |  |  |  | 5 | 6 | 6 |  |  | 1 | 1 | **19** |
| *Rudbeckia hirta var. pulcherrima* | RUDHIR |  |  |  |  |  |  |  | 2 | 61 | 4 | 88 | 7 | 32 |  | **194** |
| *Strophostyles leiosperma* | STRLEI |  |  |  |  |  |  |  | 2 | 3 |  |  |  |  |  | **5** |
| *Schizachyrium scoparium* | SCHSCO |  |  |  |  |  |  |  | 1 | 69 | 61 | 108 | 49 | 31 | 1 | **320** |
| *Lespedeza capitata* | LESCAP |  |  |  |  |  |  |  |  | 21 | 37 | 76 | 5 | 33 | 3 | **175** |
| *Rudbeckia subtomentosa* | RUDSUB |  |  |  |  |  |  |  |  | 14 | 12 | 36 | 22 | 21 | 2 | **107** |
| *Lespedeza virginica* | LESVIR |  |  |  |  |  |  |  |  | 4 | 88 | 279 | 61 | 127 | 8 | **567** |
| *Monarda fistulosa subsp. fistulosa* | MONFIS |  |  |  |  |  |  |  |  | 2 |  | 2 |  |  |  | **4** |
| *Silphium integrifolium var. integrifolium* | SILINT |  |  |  |  |  |  |  |  | 2 |  |  |  |  |  | **2** |
| *Heliopsis helianthoides* | HELHEL |  |  |  |  |  |  |  |  | 1 | 1 | 7 |  |  |  | **9** |
| *Conyza canadensis* | CONCAN |  |  |  |  |  |  |  |  | 1 |  |  |  |  |  | **1** |
| *Cyperus echinatus* | CYPECH |  |  |  |  |  |  |  |  | 1 |  |  |  |  |  | **1** |
| *Echinochloa crus-galli* | ECHCRU |  |  |  |  |  |  |  |  | 1 |  |  |  |  |  | **1** |
| *Lespedeza cuneata* | LESCUN |  |  |  |  |  |  |  |  |  | 30 | 29 | 35 | 267 | 95 | **456** |
| *Tridens flavus* | TRIFLA |  |  |  |  |  |  |  |  |  | 3 |  |  |  |  | **3** |
| *Solidago altissima* | SOLALT |  |  |  |  |  |  |  |  |  | 1 | 90 | 11 | 10 | 1 | **113** |
| *Vernonia sp.* | VERSPP |  |  |  |  |  |  |  |  |  | 1 | 1 | 1 | 1 |  | **4** |
| *Cerastium sp.* | CERSPP |  |  |  |  |  |  |  |  |  | 1 |  |  |  |  | **1** |
| *Ipomoea hederacea* | IPOHED |  |  |  |  |  |  |  |  |  | 1 |  |  |  |  | **1** |
| *Salvia azurea* | SALAZU |  |  |  |  |  |  |  |  |  | 1 |  |  |  |  | **1** |
| *Symphyotrichum sp.* | SYMSPP |  |  |  |  |  |  |  |  |  |  | 1 | 19 | 70 | 7 | **97** |
| *Gentiana sp.* | GENSPP |  |  |  |  |  |  |  |  |  |  | 1 | 1 | 21 | 2 | **25** |
| *Helianthus mollis* | HELMOL |  |  |  |  |  |  |  |  |  |  | 1 |  |  |  | **1** |
| *Coreopsis tripteris* | CORTRI |  |  |  |  |  |  |  |  |  |  |  | 2 |  |  | **2** |
| *Sporobolus compositus* | SPOCOM |  |  |  |  |  |  |  |  |  |  |  | 2 |  |  | **2** |
| *Baptisia alba var. macrophylla* | BAPALB |  |  |  |  |  |  |  |  |  |  |  | 1 | 5 |  | **6** |
| *Eupatorium sp.* | EUPSPP |  |  |  |  |  |  |  |  |  |  |  | 1 | 1 |  | **2** |
| *Eragrostis spectabilis* | ERASPE |  |  |  |  |  |  |  |  |  |  |  |  | 13 |  | **13** |
| *Pycnanthemum pilosum* | PYCPIL |  |  |  |  |  |  |  |  |  |  |  |  | 3 | 3 | **6** |
| *Boltonia asteroides* | BOLAST |  |  |  |  |  |  |  |  |  |  |  |  | 2 |  | **2** |
| *Carex bicknellii* | CARBIC |  |  |  |  |  |  |  |  |  |  |  |  | 1 |  | **1** |
| *Setaria sp.* | SETSPP |  |  |  |  |  |  |  |  |  |  |  |  |  | 1 | **1** |
| *Solidago nemoralis* | SOLNEM |  |  |  |  |  |  |  |  |  |  |  |  |  | 1 | **1** |
| **Total** | **-** | **2471** | **5479** | **452** | **318** | **96** | **150** | **232** | **121** | **559** | **743** | **1368** | **521** | **1229** | **137** | **13876** |
